# HMA proteins produce 2′,3′-cNMP signaling molecules and activate CNL-mediated immunity in rice

**DOI:** 10.64898/2026.03.15.711844

**Authors:** Xiangyu Gong, Guoqing Niu, Shuyi Sun, Bingxiao Yan, Ziqiang Li, Wei Tang, Pei Hu, Huakun Zheng, Meilian Chen, Nicolas Ponceler, Zhougeng Xu, Xing Lv, Hui Lin, Jiyun Liu, Yuanyuan Gao, Lu Zhu, Xin Wang, Guo-Liang Wang, Didier Tharreau, Houxiang Kang, Yiwen Deng, Zonghua Wang, Yu Zhang, Zuhua He

**Author notes:** These authors contributed equally (Xiangyu Gong, Guoqing Niu, Shuyi Sun, Bingxiao Yan, Ziqiang Li, Wei Tang, Pei Hu). **Correspondence:** (H.X.), (Y.W.), (Z.H.), (Y.Z.), (Z.H.).

## Abstract

Plants deploy a sensor repertoire, usually NLRs, to perceive pathogen effectors and activate executor NLR-triggered immunity. Here we reveal that rice HMA proteins function as sensors detecting the *Magnaporthe oryzae* MAX effector AvrPigm to trigger broad-spectrum blast resistance mediated by the CNL PigmR. AvrPigm is conserved with multiple copies in blast genomes and targets HMA proteins, facilitating HMA translocation into the cytoplasm. Cryo-EM structure of the HPP04^HMA^ reveals a filament-like oligomer with ssDNA/RNA bound inside the filament, which display cyclic nucleotide synthase activity to generate 2′,3′-cNMP in a CNL-dependent manner. We further discovered that the LRR domains of the executor CNLs perceive 2′,3′-cNMP to mount immunity. Our study thus establishes an unrecognized immunity mode with a distinct repertoire of sensor receptors that produce signaling molecules that are perceived by the LRR domains of executor NLRs to mediate immunity, shedding new light on the sensor-executor conception and immune activation in plants.

**Highlights:**

- HMA proteins and CNL receptors form a new sensor-executor mode in plant immunity
- Cryo-EM reveals the filament-like structure of HMA-ssRNA/ssDNA
- HMA proteins are cyclic nucleotide synthetases to generate 2′,3′-cNMP
- LRR domains of CNLs perceive 2′,3′-cNMP to mediate immunity

## INTRODUCTION

Plants have evolved a two-layer immune system to mount resistance against pathogens, which consists of pattern recognition receptor (PRR)-triggered immunity (PTI) and effector-triggered immunity (ETI) ^1,2^. ETI is usually mediated by intracellular NLR immune receptors, when activated, often induces hypersensitive responses (HR) including cell death ^3^. Based on N-terminal domains, NLRs can be divided into three classes, TNL with an N-terminal Toll/interleukin-1 receptor (TIR) domain, CNL with an N-terminal coiled-coil (CC) domain, and RNL with an N-terminal RPW8 domains ^4^. The dicot model *Arabidopsis* genome encodes both CNLs and TNLs. In contrast, the monocot model rice genome encodes CNLs and TIR-domain proteins ^5^. TNLs can form tetrameric resistosome and their TIR domains possess NAD^+^ hydrolase activity that catalyzes NAD^+^ or DNA/RNA to produce several kinds of small nucleotide molecules such as 2′cADPR, 3′cADPR, 2′,3′-cAMP/cGMP to transmit immune signal ^6–9;^ while some CNLs and RNLs form pentameric resistosomes and function as Ca^2+^ channel to trigger cell death and immune response ^10–13^. Moreover, TIR proteins can catalyze small immune molecules and promote NRG1 or ADR1-mediated immunity ^14,15^. We recently revealed that the rice TIR-only protein OsTIR also produces pRib-AMP/ADP, which trigger formation of an OsEDS1-OsPAD4-OsADR1 (EPA) immune complex ^16^. Rice also produces 2′,3′-cAMP which functions in cold tolerance and thermotolerance ^17,18^. However, there are two fundamental unanswered questions in plant immunity in monocot plants that encode only CNLs and limited TIR proteins ^16,19^: 1) how these diverse immune signals are generated, and 2) how CNLs are activated, given that the majority of CNLs do not directly perceive effectors.

The rice blast pathogen *Magnaporthe oryzae* (synonym *Pyricularia oryzae*) is one of the most destructive fungal pathogens of rice worldwide ^20^ and a model system to study plant-fungal pathogen interactions ^21^. During infection, the blast fungus secrete MAX (*M. oryzae* avirulence and ToxB-like) effectors ^22^ to suppress host immunity for pathogenesis or activate immunity upon perception by cognate NLRs encoded by blast resistance *Pi* genes ^19^. Some NLRs, such as Pik1, RGA5/Pias-1, contain integrated Heavy Metal Associated (HMA) domains ^23,24^, which bind MAX effectors to release the executor/helper NLRs’ action on immune activation ^25,26^. The HMA domain of Pik1 can be engineered to immune receptor-nanobody fusions to generate resistance against plant viruses ^27^. Moreover, HMA domain of NLRs in potato also recognize non-MAX effectors from *Phytophthora infestans* ^28^. However, how the HMA-MAX interaction releases the executor NLR action or the biochemical importance of the interaction remains undefined. In addition, the rice genomes encode 88 predicted HMA proteins, including HIPPs (heavy metal-associated isoprenylated plant proteins), HPPs (heavy-metal associated plant proteins), and other fused proteins ^29^. HIPP proteins, HIPP19 and HIPP43, are targeted by the MAX effector AvrPikD and PWL2 to suppress rice or barley immunity in the absence of the cognate NLRs ^30,31^. Both engineering HIPP19 or HIPP43 into the Pik1 HMA domain triggers Pik-mediated immunity ^32,33^. Whether these HMA proteins also serve as immune sensors directly remains an open question.

The NLR PigmR confers broad-spectrum and durable resistance to rice blast by recruiting a new transcription factor family, safeguarding primary defense metabolism and trafficking to the PM (Plasma Membrane) ^34–37^. However, the cognate effector AvrPigm and how it is perceived to trigger PigmR-mediated immunity remain unknown. Here, we identify AvrPigm through comprehensive genetic and genomic studies and find that it is recognized by specific HMA proteins, which form a filament-like structure with nucleic acids and display cyclic nucleotide synthetase (CNS) activity and produce 2′,3′-cNMP (2′,3′-cAMP/cGMP). We reveal that the LRR domain of PigmR and other allelic or paralogous NLRs bind 2′,3′-cNMP to execute immune activation. We thus reveal a previously unappreciated immunity mechanism where HMA proteins sense pathogen effectors to produce signal molecules, which are perceived by executor NLRs to activate robust disease resistance.

## RESULTS

### AvrPigm is a new MAX effector with multiple copies in *M. oryzae*

Because of the unavailability of virulent isolates to *Pigm* from the natural field (Deng et al., 2017), we identified *AvrPigm* using an integrated strategy combining artificial mutagenesis, genetic mapping, and genome sequencing. We first created virulent isolates towards *Pigm* through two continuous UV mutagenesis of the avirulent isolate TH12 (Figures S1A and S1B), obtaining TM21 ^36^ and the other 11 isolates that were fully virulent to *Pigm*, all the mutants remained virulent to Nipponbare (NIPB) (Figure S1C). In parallel, *Pigm*-virulent isolates were obtained through DEB (Diepoxybutane) mutagenesis of the avirulent isolate YN716 (Figures S1D and S1E). To genetically map *AvrPigm*, we generated 111 F1 progenies by crossing strains TM21 (virulence) and YN8773R-27 (avirulence) (Figures 1A and 1B). The segregation observed for avirulence to *Pigm* was consistent with a single locus: 55 and 56 progenies were avirulent and virulent to *Pigm*, respectively (Table S1). Through map-based cloning with bulk segregant analysis (BSA), we located a candidate locus for *AvrPigm* in a 5.42 Mb∼5.69 Mb region on Contig 3 of isolate TH12, near the telomere (Figures 1C and S1F). This region is highly enriched with repetitive sequences, precluding further fine mapping (Figure S1G). We further re-sequenced five independent virulent mutants through PacBio HiFi. Comparative genomic and collinearity analysis revealed that a large telomeric structural variation (SV) of ∼45 kb shared by virulent isolates (TM14 and TM21) but absent in wild type TH12 (Figures 1D and S1H). Within this deletion region, we identified one gene that could restore the avirulence of the mutant isolates by genetic complementation (Figure 1E). Therefore, we identified *AvrPigm*.

**Figure 1.**
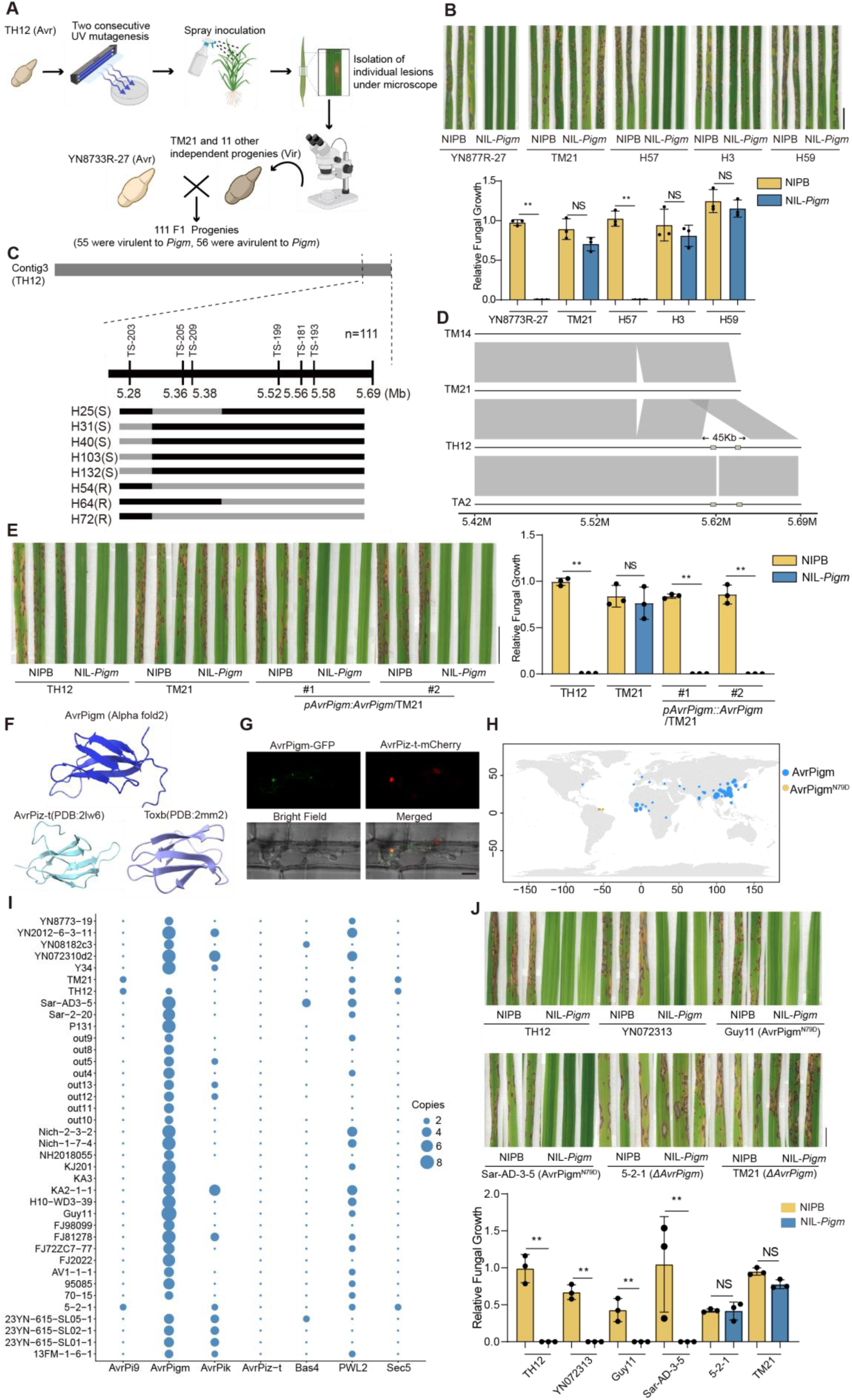
Identification and characterization of AvrPigm in *Magnaporthe oryzae*. (A) Schematic representation for genetic mapping. Two consecutive rounds of UV ray mutagenesis produced 12 independent virulent isolates to NIL-*Pigm* plants. A total of 111 F1 progenies generated by a sexual cross between isolates TM21 and YN8773R-27. Avr, avirulence. Vir, virulence. Pictures are generated by Biorender and modified in Adobe Illustrator. (B) Representative symptoms of parental and progeny isolates. Asterisks represent significant differences. NS, no significant difference. (C) *AvrPigm* was mapped to a 300kb telomeric region of Contig3 in TH12 genome. Black and gray rectangles represent respective segments from TM21 and YN8773R-27. (D) Structural variation in the *AvrPigm* region. Collinearity analysis of PacBio-resequenced TH12 (avirulence), TM21 and TM14 (virulence) revealed a large deletion in TM21. Avirulent isolate TA2 from the mutagenesis population served as a negative control. (E) Complementation of *AvrPigm* restores avirulence in TM21. The full-length *AvrPigm* sequence, including promoter and downstream regulatory regions, was transformed into TM21. Two independent transformants were sprayed on NIL-*Pigm* plants, lesions were recorded at 7 dpi. (F) Alphafold2 predicted structure of AvrPigm (blue), compared to known MAX effectors, AvrPiz-t (PDB:2lw6, indigo) and Toxb (PDB:2mm2, purple). (G) Translocation of AvrPigm into rice cells during infection. Inoculating Guy11 expressing *pAvrPigm:AvrPigm*:GFP and *pAvrPiz-t:AvrPiz-t*:mCherry on sheath cells of rice, 30 hpi. Scale bar, 10 µm. (H) The geographical distribution of AvrPigm in global populations. (I) Long-read sequencing genomes of *M. oryzae* reveals multiple copies of AvrPigm. (J) (J)Functional resilience of PigmR-mediated immunity to AvrPigm variants. AvrPigm^N79D^ (Guy ll and Sar-AD-3-5) failed to evade recognition by PigmR. natural virulent *ΔAvrPigm* (5-2-1) rendered virulence to *Pigm*. Scale bar, 1 cm (B, E, J); Error bars, mean ±SD (n = 3); **, P < 0.01 (n = 3, biological repeats), NS, no significant difference, Student’s t-test (B, E, J). See also Figure S1 and Table S1.

AvrPigm is predicted to possess a N-terminus secretion signal peptide but lacks any sequence similarity to known effectors (Figure S1I). Based on AlphaFold2 prediction ^38^, AvrPigm adopts multiple anti-parallel beta sheets, resembling MAX family effectors ^25^, including AvrPiz-t and Toxb (Figure 1F). We adopted a live-cell imaging approach ^39^ to determine the localization of AvrPigm during the infection process. AvrPigm was expressed under its native promoter with a C-terminal fusion of GFP (*pAvrPigm*:AvrPigm:GFP), and co-expressed with *pAvrPiz-t*:AvrPiz-t:mCherry, which served as a positive control. Confocal microscopy of infected rice sheaths at 30 h post-inoculation (hpi) revealed GFP fluorescence specifically localized to BICs (Biotrophic Interfacial Complex) (Figure 1G), an interfacial structure corresponding to the first hyphal cells to invade the host cells ^39^. The result confirmed that AvrPigm is secreted into BICs before delivery into the rice cell.

To elucidate the diversity of *AvrPigm* in natural *M. oryzae* populations, we analyzed ∼210 *M. oryzae* genome, including publicly available datasets ^40,41^ and newly sequenced isolates, revealed that *AvrPigm* resides in a highly repetitive genomic region and exhibits nearly identical size and coding sequence across global populations (Figure 1H and Table S1). Further long-read sequencing revealed multiple copies of *AvrPigm* (Figure 1I and Table S1), indicating that *AvrPigm* has expanded in the genomes through duplication. Only one amino acid mutation (AvrPigm^N79D^) was detected in two isolates, Guy11 and Sar-AD-3-5 from French Guyana (South America), and Suriname (South America), respectively (Figure 1H). Inoculation experiments showed that the two isolates were still avirulent to *Pigm* (Figure 1J). Moreover, we sequenced isolate 5-2-1 through PacBio HiFi, a recently naturally occurring virulent isolate from blast nursery to *Pigm*, and found a large deletion encompassing *AvrPigm* locus (Figures 1J and S1J). Therefore, we inferred that only the loss-of-function of *AvrPigm* can escape the recognition by *PigmR*. Given the characteristic feature of multi-copies, we suggest that AvrPigm is a conserved core effector and rare to lose function, providing a mechanistic explanation for the broad-spectrum and durable resistance mediated by the cognate receptor *PigmR* ^34^.

### AvrPigm targets specific HMA proteins

Yeast two-hybrid (Y2H) failed to detect a direct interaction between AvrPigm and PigmR (Figure S2A), suggesting that PigmR does not directly recognize AvrPigm. MAX effectors are frequently recognized by the integrated domain (ID) HMA of sensor NLRs, thereby triggering the activation of the executor/helper NLRs ^42,25^. However, PigmR lacks ID ^34^. Moreover, MAX effectors also target some HMA-only proteins with unclear biochemical function ^30,31^. Thus, we postulated that AvrPigm likely targets certain HMA proteins.

Using a comprehensive cereal HMA library ^29^, we annotated 88 HMA proteins in the rice genome (NIPB), and screened potential AvrPigm-interacting HMA proteins by Y2H (Figure S2B; Table S2), including HMA-only proteins and HMA domains of NLRs (e.g., Pik and RGA5). Finally, we identified four HMA proteins (HPP04, HIPP16, HIPP19, and HIPP20) that specifically interact with AvrPigm; while no interaction was observed with other HMA proteins, including Pi21 ^43^ (Figures 1A and S2B). Moreover, none of these four HMA proteins interacted directly with AvrPi9 or AvrPiz-t, highlighting MAX-HMA recognition specificity (Figure S2C). These interactions were validated independently through Co-immunoprecipitation (Co-IP) and pull-down assays (Figures 2A and 2B). Bimolecular fluorescence complementation (BiFC) assay demonstrates that AvrPigm interacts with four HMA proteins in the PM and cytoplasm (Figure 2C). Consistently, transient co-expression of GFP-tagged HIPP19 and Myc-tagged MAX effectors in *N. benthamiana* revealed that AvrPigm, but not the non-interacting AvrPi9, induced cytoplasmic relocalization of HIPP19, a similar effect was observed for HPP04 (Figures 2E, 2F and S2D). Together, these results demonstrate that AvrPigm selectively targets a set of HMAs and alters their subcellular localization, providing a mechanistic basis for PigmR-mediated immune activation.

**Figure 2.**
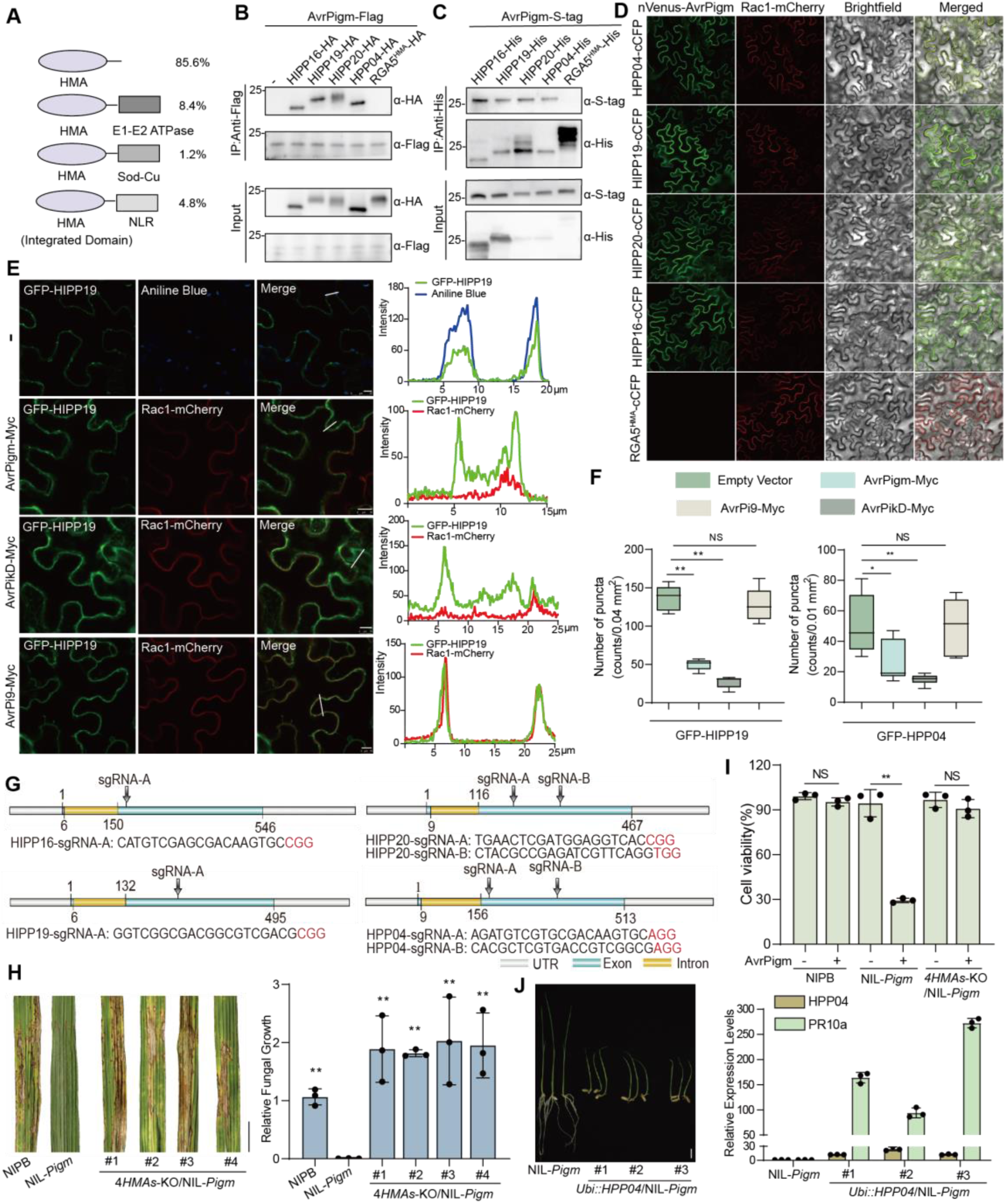
AvrPigm targets HMA proteins to activate PigmR-mediated immunity. (A) Schematic diagram of the domain organization and percentages of the rice HMA family proteins. (B) AvrPigm interacts with multiple HMA proteins in Co-IP assay. The HMA protein RGA5^HMA^ serves as a negative control. Immunoblots probed with anti-Flag/HA. (C) Pull-down assay validates the interaction between AvrPigm and four HMA proteins. RGA5^HMA^ serves as a negative control. (D) BiFc reveals AvrPigm-HMA subcellular interaction. Rac1 serves as PM marker, and RGA5^HMA^ as negative control. Scale bar, 10 µm. (E) AvrPigm alters the subcellular localization of HIPP19. N-terminal GFP-tagged HIPP19 (GFP-HIPP19) was transiently expressed in *N. benthamiana* leaves by agroinfiltration together with the indicated proteins. Aniline blue stained the plasmodesmata-associated callose. Scale bar, 10 µm. (F) Changes of GFP-HIPP19 and GFP-HPP04 puncta in the plasma membrane of *N. benthamiana*. AvrPikD serves as positive control while AvrPi9 as negative control. **, p<0.01, Student’s t-test (n = 5). (G) Sequences and genomic locations of CRISPR-Cas9 sgRNA in HPP04, HIPP16, HIPP19 and HIPP20 genes. (H) Functional redundancy of HMA proteins in NIL-*Pigm*. NIL-*Pigm* and 4*HMAs*-KO/NIL-*Pigm* lines at 7 dpi with punch injection inoculation (TH12). NIPB served as susceptible control. Different letters indicate significant differences at *P* < 0.05 (n = 3, one way-ANOVA). (I) AvrPigm-triggered cell death is compromised in the quadruple *HMA* mutants. Ectopic expression of AvrPigm in NIPB, NIL-*Pigm* and quadruple *HMA*-KO/NIL-*Pigm* protoplasts. Cell viability was measured by Titer-Glo Luminescent cell viability assay. **, p<0.01, Student’s t-test (n = 3). (J) Overexpression of HPP04 caused stunted growth in NIL-*Pigm*. Shown as morphology of seven-day-old surviving *Ubi::HPP04* seedlings in NIL-*Pigm*, and relative expression levels of *HPP04* and *PR10a* determined by qRT-PCR. NS, no significant difference (E, G), scale bar, 1 cm (H, J). See also Figure S2 and Table S2.

### HMA proteins function redundantly and induce NLR-specific autoimmunity

To define the role of HMAs in NLR-mediated immunity. We developed knockout (KO) mutants of single, double, triple, and quadruple HMAs using CRISPR-Cas9 in NIL-*Pigm* (Figures 2G, S2E and S2F). Pathogen inoculation revealed that only the quadruple KO mutants (*4HMAs*-KO, *hipp16/hipp19/hipp20/hpp04*) exhibited significantly compromised *PigmR*-mediated resistance (Figures 2H and S2G), suggesting functional redundancy of these HMA proteins. Moreover, AvrPigm triggered PigmR-mediated cell death was also abolished in the quadruple mutants (Figures 2I and S2H), demonstrating that the HMAs are required for AvrPigm-triggered PigmR activation.

To further establish the interplay between these four HMAs proteins and PigmR, we generated constitutive overexpression (OE) lines of the individual HMA proteins (HPP04, HIPP16, HIPP19, and HIPP20) in NIL-*Pigm* plants. Strikingly, most *HMA*-OE/NIL*-Pigm* seedlings were dead, while a few surviving lines exhibited stunted growth, moderate HMA expression levels and elevated expression of pathogen-related (PR) genes, and ultimately lethality (Figures 2J, S2I, and S2J). Similar autoimmunity phenotypes were also observed upon HMAs overexpression in *Pizt* (a paralogous/allelic NLR of *Pigm*) background (Figures S2K and S2L). In contrast, *HMAs*-overexpression in NIPB (without *Pigm*) showed a normal growth morphology, despite substantially higher HMA expression levels (Figure S2L). Together, these results establish that the four HMAs function redundantly and are genetically required for PigmR and other close NLR-mediated resistance.

### HMA proteins exhibit nuclease and cyclic nucleotide synthetase activity

Next, we sought to elucidate how these HMA proteins activate PigmR-mediated immunity. Structural alignment using the DALI server ^44^ revealed the similarity between monomeric structure of HMAs and *Bacillus halodurans* Cas2 (Bha_Cas2) ^45^, as well as other Cas2 homologs (Figures 3A and S3A), suggesting that the HMAs might possess an active nuclease domain. We thus performed nuclease activity assay with the above HMAs towards DNA and RNA substrates. The purified four HMAs, but not the RGA5^HMA^, displayed degrading activity towards rice genomic DNA or RNA (Figures 3B, 3C and S3B). Consistently, the nuclease activity of the HMAs was further confirmed by using a fluorescent DNA substrate (Figure 3D). Therefore, we conclude that these four HMA proteins possess bona fide nuclease activity towards both DNA and RNA.

**Figure 3.**
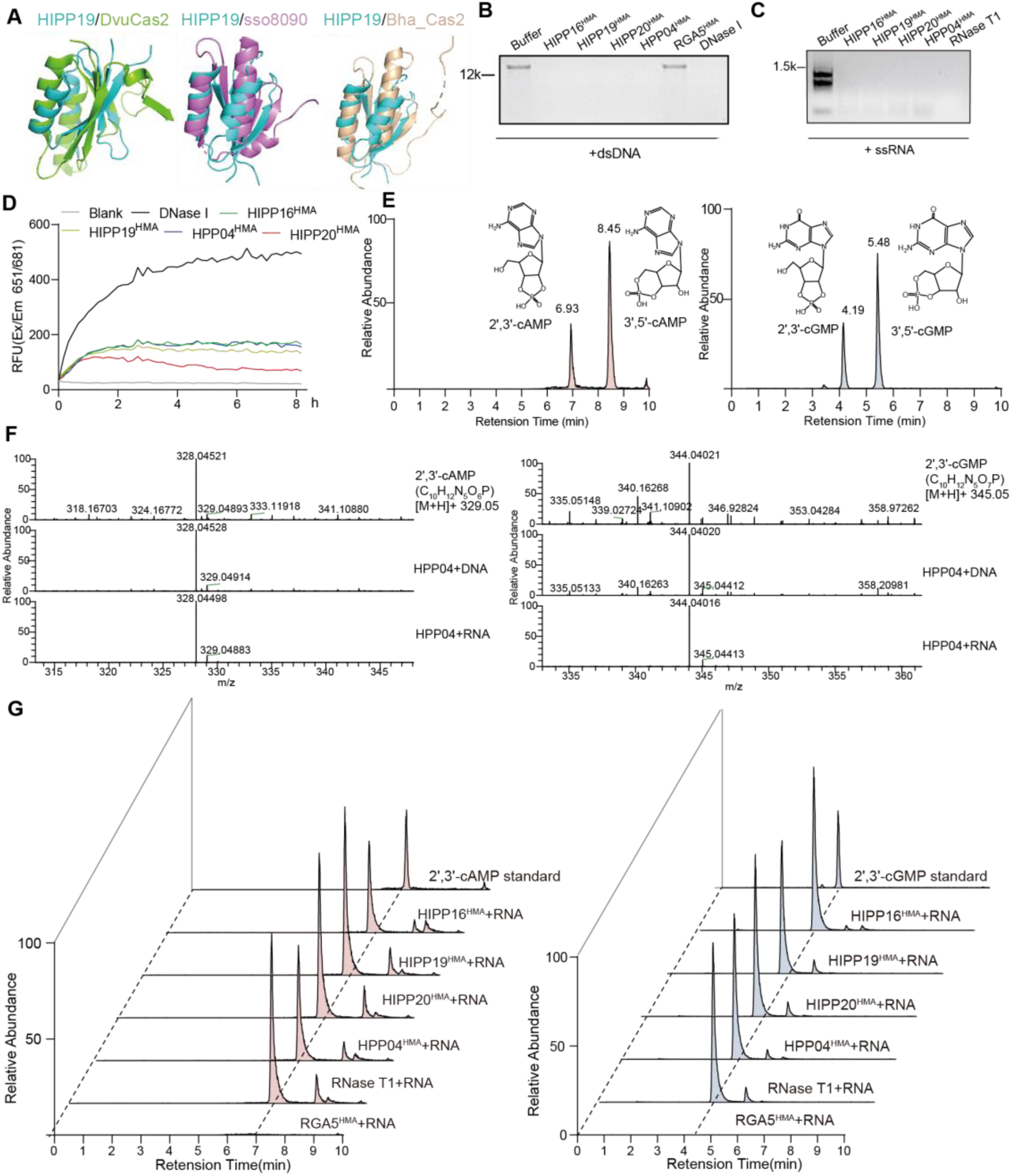
HMA proteins mimic the structure of Cas2 nuclease and exhibit both nuclease and 2′,3′-cAMP/cGMP synthetase activity. (A) Structural alignment of HIPP19^32^ with Cas2 nucleases^45^. The HMA domain of HIPP19 (PDB: 7b1i, cyan) adopts a fold similar to canonical Cas2 nucleases, including *Desulfovibrio vulgaris* Cas2 (DvuCas2, PDB: 3oq2, green; RMSD = 3.283 Å), *Sulfolobus solfataricus* Sso8090 (PDB: 3exc, purple; RMSD = 2.997 Å) and *Bacillus halodurans* Cas2 (PDB: 4es1, orange; RMSD = 3.742 Å). All structural alignments were performed using PyMOL, with water molecules and metal ions were hided. (B) DNase activity of HMA domains in vitro. Four HMA domains of (HIPP16, HIPP19, HIPP20, HPP04) can degrade rice gDNA, while RGA5^HMA^ can not. DNase І acts as positive control. Data are depicted as representative gels of three independent experiments. (C) RNase activity of HMA proteins in vitro. Proteins were incubated with rice RNA for 6 h. RNase T1 acts as positive control. Representative gels from three independent experiments are shown. (D) HMA proteins display DNase activity toward a fluorescent probe. The HMA proteins indicated were incubated with a fluorescent DNA probe. The yielded fluorescent products were measured at 646/686 nm (extinction/emission). Representative line graphs from three independent experiments are shown. (E) Molecular structures and retention time of 2′,3′-cAMP/3′,5′-cAMP and 2′,3′-cGMP/3′,5′-cGMP standards. (F) High-resolution MS spectra of the standard 2′,3′-cAMP, 2′,3′-cGMP and HMA proteins-generated products. 2′,3′-cAMP standard [M-H]^-^:328.0452, assay product [M-H]^-^:328.0449; 2′,3′-cGMP standard [M-H]^-^:344.0402, assay product [M-H]^-^:344.0401. Samples were analyzed by UPLC-Q-Orbitrap-HRMS (Thermo Fisher Scientific) and a HESI ionization source under negative ion modes. (G) LC-MS traces of 2′,3′-cAMP/cGMP from the reaction products of four HMA proteins incubating with rice RNA. HMA proteins, along with RNase T1 were incubated with rice RNA at 25°C for 6 h, and reaction products were analyzed by LC-MS relative to their standards. Chromatograms are shown for 2′,3′-cAMP (Left) and 2′,3′-cGMP (Right). Representative results from three independent experiments are shown (F, G). See also Figure S3.

We next explored the products of HMA-catalyzed degradation of DNA or RNA. Reaction product derived from RNA were analyzed by a Dionex Ultimate 3000 RSLC (HPG) ultraperformance liquid chromatography coupled with a Q Exactive quadrupole orbitrap high-resolution mass spectrometry (UPLC-Q-Orbitrap-HRMS). The results showed that the HMA-catalyzed RNA products exhibited retention times and masses (328 and 344 m/z) identical to those of the standard cyclic nucleotides 2′,3′-cAMP and 2′,3′-cGMP, with RNase T1 serving a reaction control (Figures 3E-3G, and S3C). Similar cyclic nucleotides were detected when genomic DNA was used as the substrates, comparable to those generated by the control L7^TIR^ (Figure S3D) ^9^. Therefore, we discovered that these HMAs function as noncanonical cyclic nucleotide synthetase, catalyzing the production of 2′,3′-cAMP/cGMP with DNA or RNA as substrates.

### Cryo-EM structure reveals DNA/RNA-induced filament-like oligomerization of HPP04^HMA^

To understand the structural basis of the HMA enzymatic activity, we sought to determine the structure of HMA. During protein purification, a large proportion of HMAs was always eluted in fractions close to the void volume on a Superose 6 gel filtration column, albeit its theoretical molecular weight of 8 kDa, suggesting the formation of high-molecular-weight oligomers (Figure 4A). Notably, the oligomers exhibited significant absorbance at 260 nm (A260), indicating the presence of nucleic acids (Figure S4A). Further analysis of the HPP04^HMA^ oligomers on a negative staining electron microscopy (EM) revealed filament-like structures (Figure 4B). Similar filament-like structures were also obtained with HIPP16^HMA^, HIPP19^HMA^, and HIPP20^HMA^ (Figures S4B), suggesting a conserved filament-forming feature of the HMAs.

**Figure 4.**
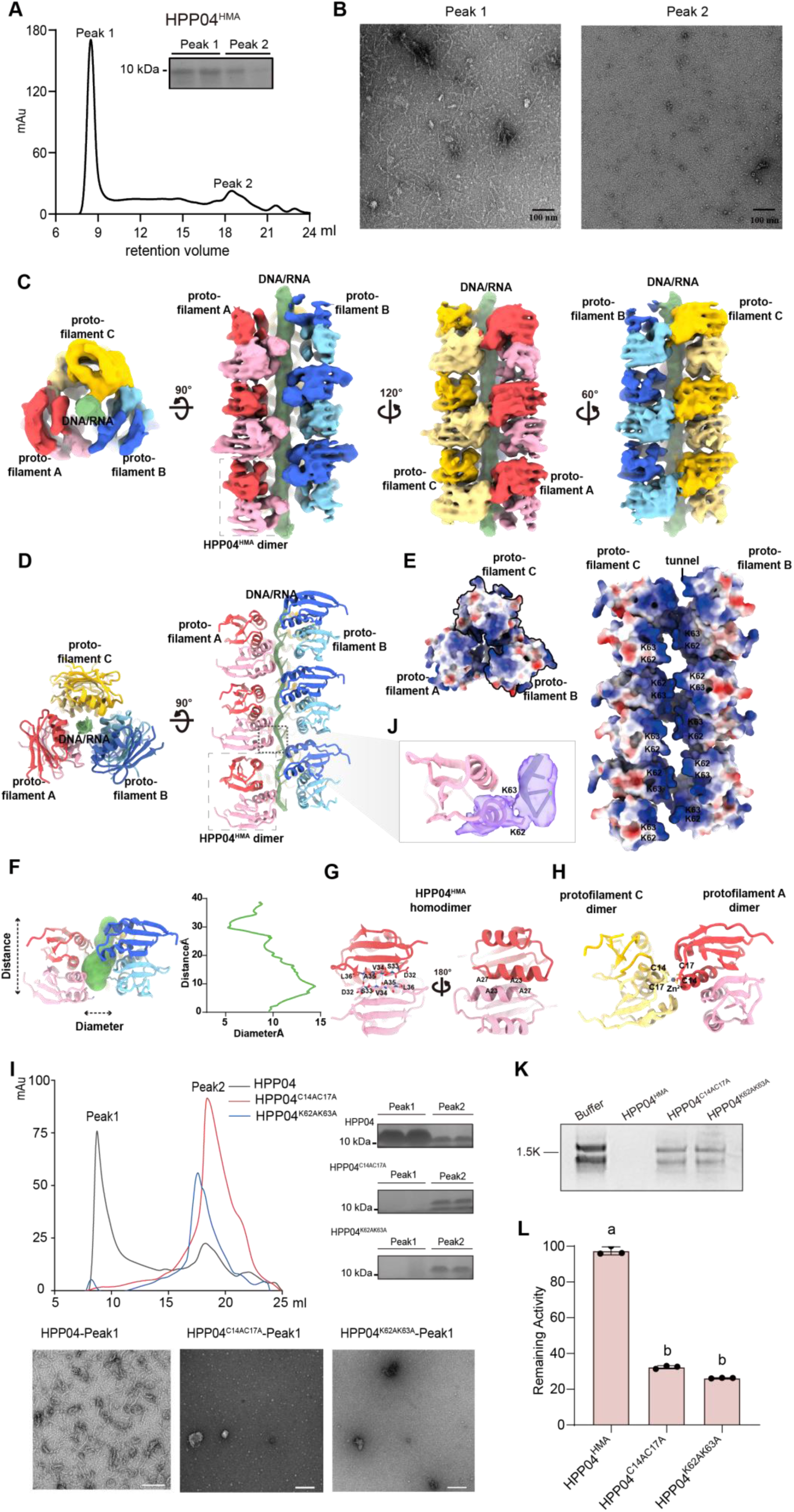
Cryo-EM structure of the HPP04^HMA^ with ssRNA/ssDNA. (A) Gel-filtration chromatography from the Superose 6 column, showing that HPP04^HMA^ exhibits two oligomerization states, peak one is eluted at 9 ml, and peak two at 19 ml. Horizontal axis: retention time (min); vertical axis: UV absorbance at 280 nm. Data are depicted as a representative result of three independent experiments. (B) Negative-staining result shows the different shapes of HPP04^HMA^. Left: HPP04^HMA^ from peak 1; right, HPP04^HMA^ from peak 2. Scale bar, 100 nm. (C) Top and side view orientations of the cryo-EM map of HPP04^HMA^ filament. (D) Top and side view orientations of the model of HPP04^HMA^ filament. (E) Top and side view orientations of the electrostatic potential map of the HPP04^HMA^ filament structure. (F) The diameter profile of the central pore, calculated using MOLEonline (Pravda et al., 2018), reveals a maximum diameter of 14.5 Å. This dimension is insufficient to accommodate double-stranded nucleic acids, which typically require diameters of 25.5 Å (A-form), 23.7 Å (B-form), or 18.4 Å (Z-form). (G) The protomers are glued together in a head-to-tail manner via an interprotomer β-sheet and a coiled coil. (H) The inter-protofilament zinc fingers formed by residues Cys14 and Cys17 facilitate assembly of the three-strand filament. (I) Mutations of Cys14 and Cys17 or Lys62 and Lys63 disrupted the HPP04^HMA^ filament. (J) Cryo-EM density of Lys62 and Lys 63 within the HPP04^HMA^-DNA/RNA filament. (K) Effect of HPP04^HMA^ mutation on nuclease activity towards RNA. (L) Effect of HPP04^HMA^ mutation on 2′,3′-cNMP synthetase activity. Different letters indicate significant differences at P < 0.05 (n = 3, one way-ANOVA). See also Figure S4, S5 and Table S3.

To elucidate the structural details of the HMA filaments, we determined the cryo-EM structure of HPP04^HMA^ filaments at a resolution of 3.54 Å (Figure 4C; Video S1), reminiscent of ‘the Tree of Life or Bronze Sacred Tree (Shang Dynasty)’. The cryo-EM map reveals that the HPP04^HMA^ filament is composed of three protein protofilaments (protofilaments A, B, and C), creating a tunnel inside with a diameter of 14.5 Å (Figure 4D). Notably, we observed a continuous, strong cryo-EM map in the middle of the tunnel, forming the fourth protofilament. We attribute the fourth filament to single-stranded DNA or RNA based on the following evidence: 1) the electrostatic analysis shows a strong positively charged potential of the inner tunnel wall, complementary to the negatively-charged phosphate backbone of a DNA/RNA strand (Figure 4E); 2) the high OD_260_/_280_ ratio of the HPP04^HMA^ sample suggests the presence of nucleic acids (Figure S4A); 3) the diameter of the tunnel cannot accommodate a double-stranded DNA (dsDNA), double-stranded RNA (dsRNA), or DNA-RNA hybrid due to steric hinderance (Figure 4F). The results of electrophoretic mobility shift assay (EMSA) using the non-oligomer form of HPP04^HMA^ showed that it could bind to various types of nucleic acids, with a preference for ssRNA (Figures S4C and S4D). We also observed a discontinuous map in the middle tunnel in a lower-resolution map reconstructed from a subset of 3D classes, which might be indicative of DNA/RNA cleavage (Figure S5)

The building unit of the HPP04^HMA^-ssDNA/RNA filament is the HPP04^HMA^ dimer, in which two protomers are glued together in a head-to-tail manner through the formation of an inter-protomer β sheet and a coiled coil (Figure 4G). The HPP04^HMA^ dimer docks with each other and forms a HPP04^HMA^ protofilament. Intriguingly, we identified an NTP molecule at the interface of two HPP04^HMA^ dimer building unit (Figure S4E). Our filaments assembly experiment shows that addition of 1 mM ATP dramatically enhanced the formation of HPP04^HMA^ filaments (Figure S4F), suggesting that ATP may act as a cofactor to stabilize the HMA protofilament. The three protofilaments are further assembled together through inter-protofilament zinc fingers formed by residues Cys14 and Cys17 (Figure 4H). Mutating these two cysteines disrupted the ability to form filaments (Figure 4I). In the middle tunnel, a lysine pair (Lys62 and Lys63) on the α2 helix of each HPP04^HMA^ protomer positions its side chains toward the central ssDNA/RNA, creating a highly positively charged electrostatic potential for interacting with the negatively charged DNA or RNA (Figure 4J). To further investigate whether these lysine residues play the structural role for HPP04^HMA^-DNA/RNA protofilament assembly, we engineered a double alanine substitution mutant (2KA, K62A/K63A). In contrast to HPP04^HMA^, the mutant protein eluted exclusively as a low-molecular-weight species in gel filtration, indicating a complete disruption of filament assembly (Figure 4I). This finding establishes that the interaction between α2-helix and nucleic acids is essential for filament formation. Strikingly, disruption of filament formation substantially impaired the enzymatic activity of HPP04^HMA^, with a significant loss of nuclease activity towards RNA substrates (Figure 4K) and a 60%-80% reduction in 2′,3′-cNMP synthetase activity (Figure 4L), suggesting that the ssDNA/RNA-induced filament formation promotes HMA enzymatic activity. In summary, HPP04^HMA^ forms a filamentous structure enclosing a ssDNA/RNA within the middle tunnel.

Collectively, we show that HPP04^HMA^ filament is formed from its dimer building blocks and requires interactions between the lysine residues of the dimer block and the enclosed ssDNA/RNA. The HPP04^HMA^ filament degrades nucleic acids and synthesizes cyclic nucleotide monophosphates.

### 2′,3′-cNMP synthetase activity of HMAs is required and promoted by PigmR

Given the autoimmune phenotype of HMAs-overexpressing in NIL-*Pigm* (Figures 2J and S2I), we further performed transient expression assays to monitor cell death in rice protoplasts. Transient expression of either HMA proteins triggered severe cell death in NIL-*Pigm* protoplasts, accompanied by significant elevated 2′,3′-cNMP levels (Figure 5A, 5B, and S6C). In contrast, HMAs expression neither triggered cell death nor significant elevated 2’,3’-cNMP levels in NIPB protoplasts (Figures S6A-S6C). To investigate whether 2’,3’-cNMP directly activate immune response, we exogenously supplied 1 mM 2’,3’-cNMP to NIPB protoplasts. No obvious cell death was observed (Figures S6D). However, in PigmR-expressing protoplasts, 2’,3’-cNMP recapitulated the cell death feature seen with HPP04-PigmR co-expression (Figures S6D). Together, these results establish that 2’,3’-cNMP functions as an immune signal in an NLR-dependent manner, functionally linking effector perception by HPP04 to immune response mediated by PigmR.

**Figure 5.**
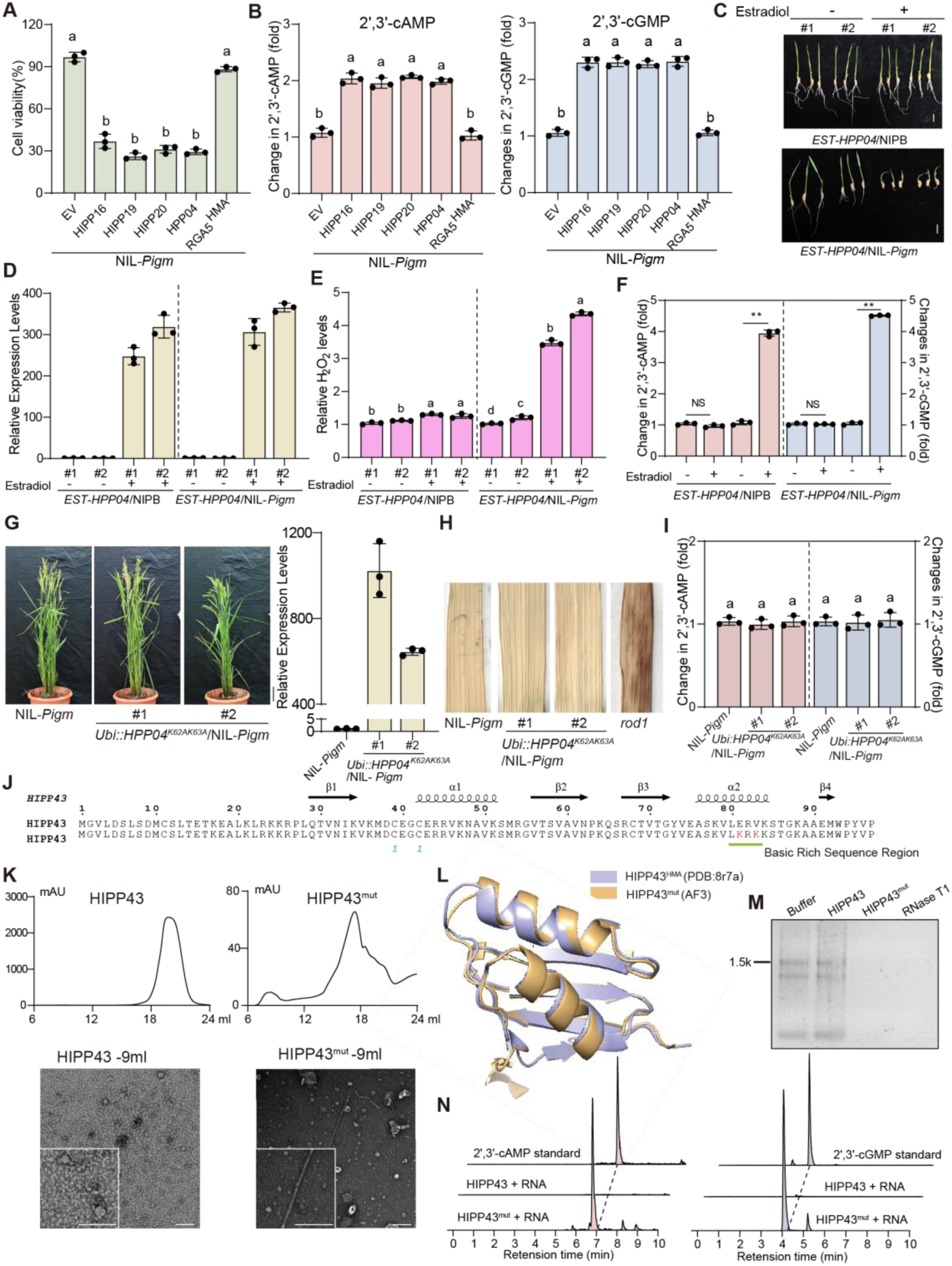
Overexpression of HMAs trigger cell death accompanied with increased levels of 2′,3′-cNMP in NIL-*Pigm* and engineering non-filamentous HMA into filamentous HMA. (A) Four HMA proteins induced strong cell death in NIL-*Pigm* protoplasts, compared to RGA5^HMA^ and empty vector (EV). Assays were performed as described for Fig.2G. (B) 2′,3′-cNMP levels in NIL-*Pigm* protoplasts expressing five HMA proteins and empty vector. The extracts were analyzed by LC-MS. Data are normalized to empty vector (EV). (C)-(F) Morphology of seven-day-old seedlings in NIPB and NIL-*Pigm* with or without 50 µM estradiol treatment (C). Relative expression levels of *HPP04* in NIPB and NIL-*Pigm* (D). Relative H_2_O_2_ levels in *EST-HPP04*/NIPB and *EST-HPP04*/NIL-*Pigm* (E). 2′,3′-cNMP levels in *EST-HPP04*/NIPB and *EST-HPP04*/NIL-*Pigm* (F). For each assay, values in DMSO-treated plants were normalized to 1. Dash lines separate the experimental groups, performing statistical analysis independently. (G) HPP04^K62AK63A^ lost autoimmunity phenotype in NIL-*Pigm*. The relative expression levels of HPP04^K62AK63A^ in Adult NIL-*Pigm* plants are shown. Scale bar,10 cm. (H) DAB staining indicating H_2_O_2_ accumulation in HPP04 ^K62AK63A^/NIL-*Pigm* l by (H). The autoimmune mutant *rod1* serves as positive control^58^. (I) 2′,3′-cNMP levels in HPP04 ^K62AK63A^/NIL-*Pigm* lines. Statistical analyses of 2′,3′-cAMP/cGMP were performed independently. (J) Engineered HIPP43 with nucleic acid-binding residues (red). (K) HIPP43^mut^ obtains the activity to form filaments in vitro. Top panels, gel-filtration chromatography profiles display the protein states of HIPP43 and HIPP43^mut^. Bottom panels, negative-staining images of HIPP43^HMA^ and HIPP43^mut^ eluted at the void position. (L) Unchanged monomeric structure of HIPP43^mut^ compared with HIPP43. Purple, HIPP43 (PDB:8r7a); orange, HIPP43^mut^ (AF3). (M) Nuclease activity for HIPP43 and HIPP43^mut^ towards RNA. (N) 2′,3′-cNMP synthetase activity of HIPP43^mut^ towards RNA. Statistical analyses were tested using Student’s t-test (* p < 0.05) (n = 3, F) or one-way ANOVA (p < 0.05, n = 3, indicated by different letters, A, B, E, I). See also Figure S6.

Further, to confirm *bona fide* HMA-driven 2′,3′-cNMP production in *planta*, we analyzed estradiol-inducible *EST:HPP04* transgenic lines. Upon 50 μM estradiol induction, *EST:HPP04/*NIL*-Pigm* seedlings exhibited stunted growth, H_2_O_2_ accumulation, and higher 2’,3’-cNMP accumulation in *EST:HPP04/*NIL*-Pigm* but not in *EST:HPP04/*NIPB plants (Figures 5C–5F). These results demonstrate that the production of 2’,3’-cNMP by HMAs requires the presence of PigmR in *planta*.

We next determine whether the nucleic acid binding ability of HMAs is essential for ETI activation. Through transgenic approaches, we found that overexpression of HPP04 nucleic acid binding (K62A/K63A) mutants in NIL-*Pigm* yielded viable plants without autoimmunity phenotypes and increased 2’,3’-cNMP levels (Figures 5G-5I), contrasting with the autoimmunity observed upon overexpression of wild type HPP04 (Figure 2J). Collectively, these results provide genetic evidence that the 2’,3’-cNMP synthetase activity of HPP04 is indispensable for NLR-triggered immunity.

Next, we observed that expressing AvrPigm in NIL-*Pigm* caused visible cell death with increased 2’,3’-cNMP levels while expressing AvrPigm in NIPB had no significant effect (Figures S6E and S6F). To further elucidate the role of 2’,3’-cNMP during the infection process, we inoculated NIL-*Pigm* plants TH12 (avirulent), TM21 (virulent), and water as a control, and subsequently quantified 2’,3’-cNMP levels. Notably, inoculation with TH12 but not TM21 resulted in the significant accumulation of 2’,3’-cNMP at 24 hpi, which returned to base levels at later time points (Figures S6G). Thus, we conclude that AvrPigm promotes the 2′,3′-cAMP/cGMP production in NIL-*Pigm,* which in turn triggers PigmR-mediated immunity.

### Engineering non-filamentous HMA into filamentous HMA to obtain cyclic nucleotide synthetase activity

To understand the sequence determinants underlying HMA filamentation, we compared the sequence of four identified filament-forming HMA proteins (HIPP16, HIPP19, HIPP20, and HPP04) (Figure S6H). Integrating insights from the cryo-EM structure, we identified two critical motifs in α helix that are essential for filament-forming: 1) a ‘CXXC (S)’ motif in α1 helix mediating protofilament interwinding; 2) a conserved LRK (R)K motif in α2 helix for interacting nucleic acid. In contrast, HMAs lacking these two critical motifs, such as Pi21^HMA^ and RGA5^HMA^, failed to form filaments and lack 2′,3′-cNMP synthetase activity (Figure S6I-S6K). Collectively, these findings suggest that rice HMA family could be classified into filamentous and non-filamentous subgroups.

Next, we sought to determine whether we could create a filamentous HMA protein by replacing the critical residues in a non-filamentous HMA and test it for nucleic acid binding gain-of-function. One HMA protein, HIPP43, whose sequence lacks the lysine pair for nucleic acid binding ^33^, failed to form filaments (Figures 5J and 5K). Replacing the α2 residues of HIPP43 with lysine-rich containing sequences resulted in modified HIPP43^mut^ (E81KV83K), which could assemble into filaments *in vitro* without altering the predicted monomeric structure (Figures 5J-5L). Moreover, HIPP43^mut^ acquired both nuclease and 2’,3’-cNMP synthetase activities (Figures 5M and 5N). These results establish that local sequence motifs rewiring is able to convert a non-filamentous HMA into a functional filamentous one with 2’,3’-cNMP production, exhibiting potential for engineering new immunity-triggering HMA proteins.

### Cytoplasmic HMAs activate PigmR-mediated immunity

To delineate the dynamics of PigmR-triggered immunity, we performed time-lapse live-cell imaging of rice protoplasts within 6-9 h post-transfection ^10^. NIL-*Pigm* protoplasts expressing HIPP19 or HPP04 were stained with SYTOX Blue at 6 h, a dye that binds to the DNA/RNA of dying cells, resulting in blue fluorescence signals, which were acquired continuously (every 30 s) for 20-60 min. Dying protoplasts were suddenly disrupted (Figures 6A and S7A; Video S2 and S3), reminiscent of canonical cell death occurrence^10^. The SYTOX Blue fluorescence signal initiated from the cytoplasm and intensified prior to cell rupture (Figures 6A and S7A), indicating the intracellular nucleic acid accumulation and cytoplasmic origin for PigmR-triggered immunity, likely through PM-associated NLR complex activation ^37^.

**Figure 6.**
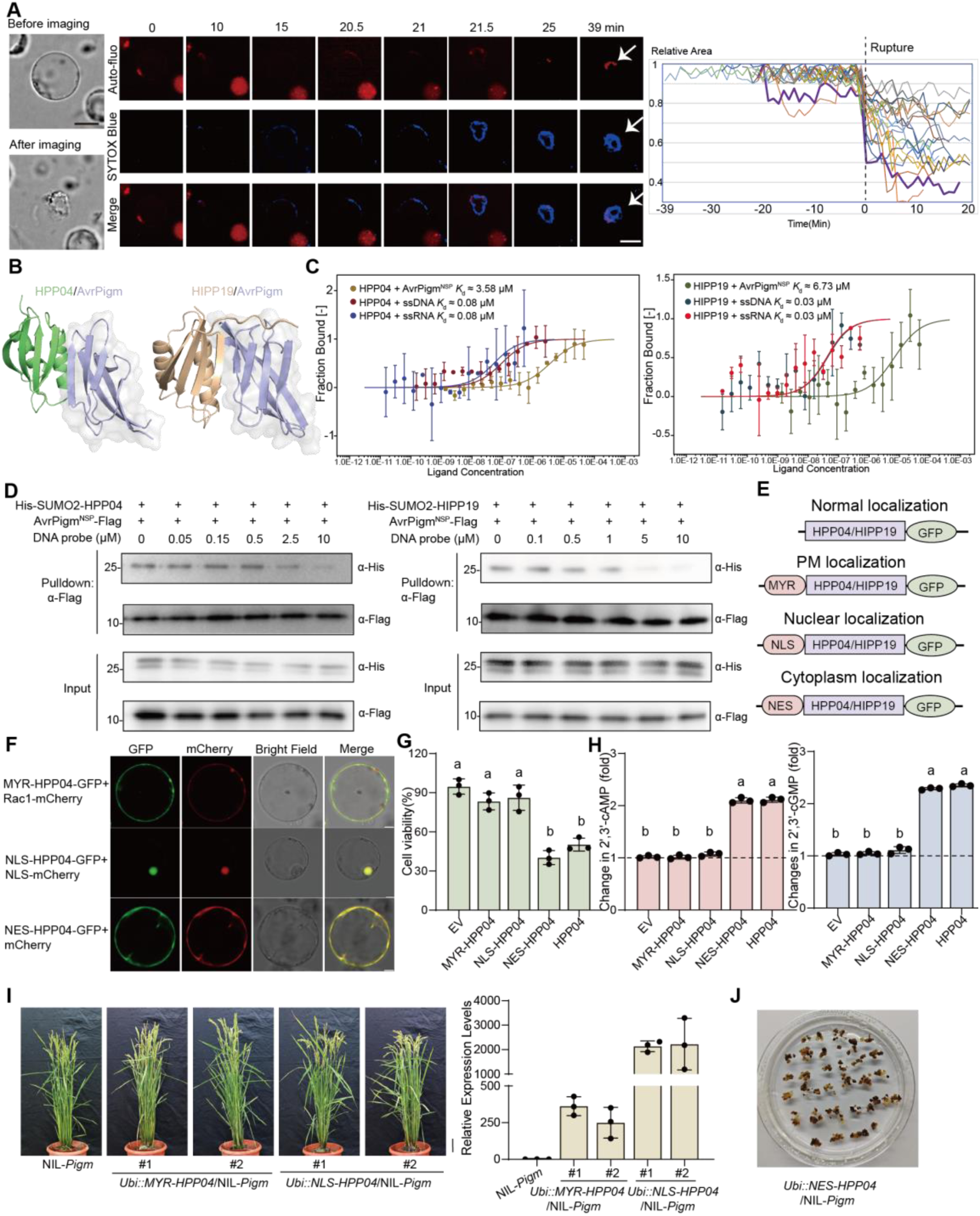
Cytoplasm-localized HMAs trigger PigmR-mediated cell death via 2′,3′-cNMP production. (A) Time-lapse imaging of HIPP19-triggered cell death in NIL-*Pigm* protoplasts, stained with SYTOX Blue 6 h post-transfection. SYTOX Blue fluorescence and chloroplast auto-fluorescence were captured every 30 s by spinning-disk microscopy. Arrowheads indicate dying protoplasts (turned SYTOX Blue). Cross sectional areas of dying protoplasts were measured and plotted against time. Rupture time was set as 0, and areas were normalized to the initial area to track trajectories (n =18). (B) Predicted complex structure of AvrPigm-HPP04 and AvrPigm-HIPP19 by AF3. (C) Quantification of the binding affinity among AvrPigm, HPP04, HIPP19 and ssDNA/ssRNA by MST. HPP04 (left) and HIPP19 (right) were labeled with Red-NHS and measured the binding affinity with the indicated protein or nucleic acids. (D) Competitive assays of AvrPigm-HPP04 and HIPP19 interactions in the presence of ssDNA. AvrPigm^NSP^-Flag (0.5 µM) was incubated with 0.5 µM HPP04-/HIPP19-His-SUMO2 for 30 min. The mixture was flowed through Flag beads with increasing ssDNA concentrations (0, 0.05, 0.15, 0.5, 2.5 and 10 µM), the bound proteins were analyzed by SDS-PAGE. Three independent experiments yielded similar results. (E) Schematic representation of subcellular localization-modified HMA fusions. HPP04/HIPP19 were fused to compartmentalization signals in N-terminus, MYR for plasma membrane; NLS (nuclear localization signal) for nucleus; NES (nuclear export signal) for cytoplasm, and C-terminal fusion to GFP for observations. (F) Subcellular localization of MYR/NLS/NES-HPP04-GFP in NIL-*Pigm* protoplasts. Showing altered subcellular location of the fusion proteins. Markers, Rac1-mCherry (PM), NLS-mCherry (nuclear), and mCherry (cytoplasm). Scale bar, 5 µm. (G) Subcellular localization modulates HPP04-induced cell death in NIL-*Pigm*. Cell viability in protoplasts expressing MYR/NLS/NES-HPP04-GFP were measured by Cell Viability Assay. Note that only cytoplasm-located fusion proteins induced cell death, compared to the HPP04-GFP. (H) Subcellular localization affects the ability of HPP04 to produce 2′,3′-cNMP. NIL-*Pigm* protoplasts expressing the indicated constructs were lysed and measured 2′,3′-cNMP levels by LC-MS. (I) Morphology of transgenic rice seedlings overexpressing MYR-HPP04 and NLS-HPP04 fusions. *Ubi::MYR-HPP04* and *Ubi::NLS-HPP04* plants exhibit no autoimmunity phenotype. Relative expression levels of HPP04 were detected by qRT-PCR. Scale bar, 10 cm. (J) Cell death of transgenic rice callus expressing NES-HPP04 in NIL-*Pigm*. Data are mean ±SD of three biological replicates. Different letters indicate significant differences at P < 0.05 (n=3, one way-ANOVA). See also Figure S7.

Given that AvrPigm interacts with HMAs and HMAs are capable of forming filaments with nucleic acids (Figures 2D, 4C and 6B), we next sought to elucidate the relationship between AvrPigm-HMA interaction and HMA binding nucleic acids in the cytoplasm. We first quantify the binding affinities of HMA proteins with AvrPigm or nucleic acids. We labelled HPP04/HIPP19 and perform Microscale thermophoresis (MST) assay, and observed that HPP04 and HIPP19 bound to AvrPigm with dissociation constants (*K*d) of approximately 3.58 μM and 6.73 μM, respectively (Figure 6C). Notably, both HPP04 and HIPP19 exhibited much stronger binding affinities to ssDNA/RNA, with *K*d values of about 0.08 μM and 0.03 μM, respectively, nearly 100-fold higher than their affinity for AvrPigm (Figure 6C). Pull-down assays further demonstrated that nucleic acid efficiently disrupted the interactions between AvrPigm and HPP04/HIPP19 (Figure 6D). Thus, we conclude that nucleic acids disrupt the AvrPigm-HMA interactions, thereby promoting HMA enzymatic activity to generate immune molecules in the presence of the cognate NLRs.

GFP fusions of C-terminal or N-terminal of HPP04 and HIPP19 revealed subcellular location to the PM, cytoplasm and nucleus in rice protoplasts (Figures S7B and S7C). To determine whether the subcellular localization of HPP04 and HIPP19 modulates PigmR-mediated cell death, we fused HPP04 and HIPP19 using targeting signal peptide for plasma membrane (MYR, myristoylation and palmitoylation sequence), nucleus (NLS, nuclear localization signal) or for cytoplasm (NES, nuclear export signal) (Figures 6E, 6F and S6D). Only NES-HPP04/HIPP19 triggered robust cell death in NIL-*Pigm* protoplasts, while MYR-HPP04/HIPP19 and NLS-HPP04/HIPP19 failed to do so (Figures 6G and S6E). Elevated 2’,3’-cNMP levels in NES-tagged samples were comparable to those observed in samples expressing wild type HPP04/HIPP19 (Figures 6H and S6F).

To further determine the compartmentalization requirement of HMA activity in *planta*, we generated transgenic NIL-*Pigm* plants expressing MYR/NLS/NES-HPP04. Interestingly, plants expressing MYR-HPP04 or NLS-HPP04 remained viable with no excessive H_2_O_2_ accumulation, and showed basal 2’,3’-cNMP levels comparable to NIL-*Pigm* controls (Figure 6I, S7G-S7I). On the contrary, cytoplasm-localized NES-HPP04/HIPP19 exhibited strong cell death in callus stage (Figures 6J and S7J), and surviving seedlings only exhibited low expression levels of HPP04/HIPP19 (Figure S7K). This phenotype recapitulates the autoimmune death caused by wild-type HPP04 overexpression in NIL-*Pigm* (Figure 2H). Given that AvrPigm promotes cytoplasmic redistribution of HMAs (Figures 2D and 2E), we propose that cytoplasmic localization of HMAs is essential for 2’,3’-cNMP production and PigmR-mediated immunity.

### 2’,3’-cNMP is perceived by LRR domains of CNLs in rice

LRR domains of PRRs are known to recognize ligands, such as FLS2-flg22 and EFR-elf18 ^46,47^. Analogously, LRR domains of some NLRs can perceive the cognate effectors or guardee proteins to activate immunity ^12,48^. A recent work also identified inositol binding to the LRR domain of SlNRC2 to mediate cell death ^49^. Since PigmR does not interact with AvrPigm directly (Figure S2A), we hypothesized that its LRR domain might recognize the 2’,3’-cNMP produced by the sensor HMAs. To address this hypothesis, we purified the LRR domain of PigmR and performed MST assay (Figures S8A and 7A). MST results suggested the LRR domain of PigmR exhibited micromolar binding affinity for both 2’,3’-cAMP (*K*d ≈ 5.2 μM) and 2’,3’-cGMP (*K*d ≈ 0.3 μM), while the LRR domain of the non-responsive CNL Pish showed much weaker binding ability (*K*d ≈ 7.9 mM and 0.3 mM, respectively) (Figure 7A). Further flow-induced dispersion analysis (FIDA), a label- and immobilization-free technology, also confirmed these interactions (Figure S8B). Similarly, paralogous CNLs Pi9 and Piz-t bound 2’,3’-cNMP via their LRR domains (Figure 7A and S8B), suggesting a conserved immune recognition mechanism of this CNL subfamily.

**Figure 7.**
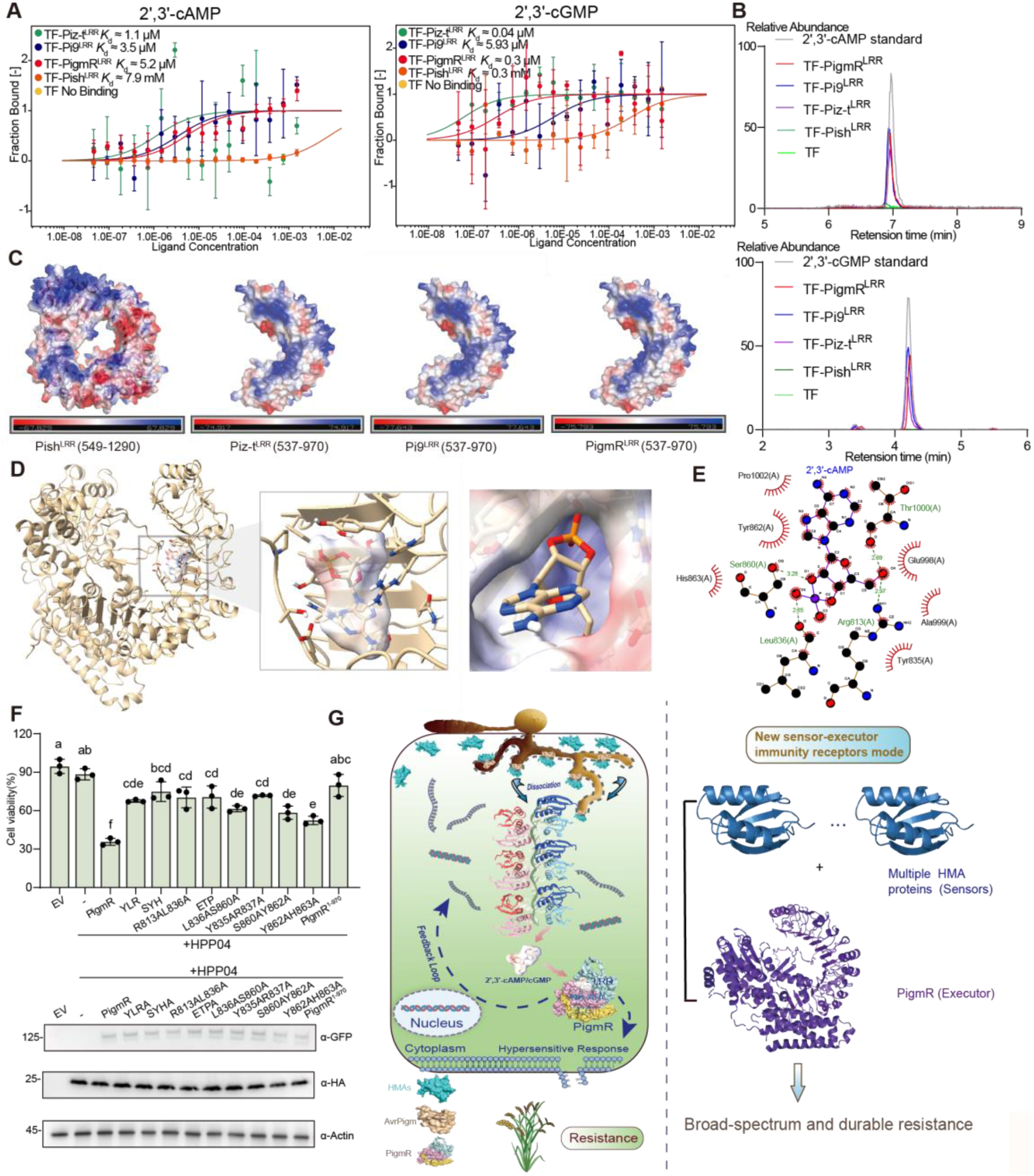
PigmR and other paralog NLRs perceive 2′,3′-cNMP via LRR domain. (A) MST quantifies binding affinity of the LRR domain of PigmR/Piz-t/Pi9/Pish to 2′,3′-cNMP. LRR domains of PigmR/Piz-t/Pi9/Pish were labeled with RED-NHS fluorescent dye and mixed with serial dilution of 2′,3′-cNMP as indicated. Note that the race-specific NLR Pish exhibited much lower (millimolar) binding affinity towards 2′,3′-cNMP. TF was used as a negative control. (B) Native MS analysis validates direct interaction between LRR domains of NLRs and 2′,3′-cNMP. LC-MS chromatograms reveals the 2′,3′-cNMP in the PigmR/Piz-t/Pi9 LRR domain complexes, while 2′,3′-cNMP was hardly detected with the Pish LRR domain. (C) The electrostatic potential distribution of Pish^LRR^, PigmR^LRR^, Pi9^LRR^, and Piz-t^LRR^. Surface analysis was performed using Pymol. Positive charge is indicated in blue and negative charge in red. (D) Predicted structure of the PigmR-2′,3′-cAMP complex by AF3. 2′,3′-cAMP is buried in the pocket of PigmR LRR domain. Electrostatic surface analysis of PigmR^LRR^-2′,3′-cAMP binding pocket. (E) Identification of critical residues at 2′,3′-cAMP-PigmR interaction interface through LigPlot. Red, predicted hydrophobic interaction, green, hydrogen bond interaction. (F) Cell viability of PigmR and its 2′,3′-cAMP-binding mutants. Mutants, YLR (PigmR^Y835AL836AR837A^), SYH(PigmR^S860AY862AH863A^), R813AL836A (PigmR^R813AL836A^), ETP (PigmR^E998AT1000AP1002A^), L836AS860A (PigmR^L836AS860A^), Y835AR837A (PigmR^Y835AR837A^), S860AY862A (PigmR^S860AY862A^), Y862AH863A (PigmR^Y862AH863A^) and PigmR^1-970^ (Deleting 971-1032). Immunoblot analysis of expression levels of the constructs. Rice actin acts as loading control. (G) A proposed model for the AvrPigm-HMAs-PigmR module in rice broad-spectrum blast resistance. Left, the conserved MAX effector AvrPigm interacts with specific HMA proteins to activate immunity. Upon infection, AvrPigm promotes cytoplasmic translocation of HMA proteins, which then assemble into filamentous structures with nucleic acids and produce 2’,3’-cNMP. The resulting 2’,3’-cNMP molecules are sensed by the LRR domain of PigmR that activate PigmR-mediated robust defense, ultimately cell rupture. Right, a new sensor-executor mode for immune activation in plants. See also Figure S8.

To further validate the specificity of CNL-LRR domain interaction with 2’,3’-cNMP, we performed native mass spectrometry (MS) on purified LRR domains of PigmR, Pi9, Piz-t and Pish. After denaturing the proteins and extracting small-molecule using a previously reported procedure ^49^, the extractants were then subjected to liquid chromatography MS (LC-MS) analysis. This process revealed that 2’,3’-cNMP was exclusively detected in the supernatants from PigmR, Pi9 and Piz-t LRR domain samples, with less 2’,3’-cNMP detected in the Pish samples (Figure 7B). Taken together, these results indicate that the LRR domains of specific NLRs bind to 2’,3’-cNMP directly, providing new evidence for how CNL LRR domains perceive immune signals and activate broad-spectrum disease resistance.

### 2’,3’-cAMP-binding is required for NLR-mediated cell death

To define the structural basis of 2’,3’-cAMP recognition by PigmR and other NLRs, structural predictions of PigmR and paralogous CNLs were carried out using Alphafold-2 ^38^. Structural alignment revealed high similarity among PigmR, Pi9 and Piz-t (RMSD = 0.603, 0.695, respectively), while Pish diverged significantly (RMSD = 8.408) (Figure S8C). Surface electrostatic potential also showed a shared ligand-binding landscape among PigmR, Pi9 and Piz-t, but not Pish (Figure 7C). We then used Alphafold-3^50^ and molecular dynamics simulations to model the 2’,3’-cAMP-PigmR complex and identified a potential ligand-binding pocket (Figures 7D, 7E and S8D). Within this pocket in the LRR domain, PigmR encodes two aromatic residues, Y862 and Y835, which stack against the nucleobase and ribose moiety of the ligand, respectively (Figure 7E). These residues are highly conserved across broad-spectrum blast resistance CNLs (e.g., Pi2, Pi9 and Piz-t), whereas they exhibit sequence variability in other rice CNLs (e.g., Pish and OsADR1) (Figure S8E). Besides the aromatic residues, the polar residues (S860, R813 and T1000) and residue L836 form hydrogen bands with the phosphate and ribose hydroxyl groups (Figure 7E). Functional mutagenesis of key pocket residues, as well as disruption of adjacent β-strand contact regions (Y835-L836-R837 and S860-Y862-H863) or the hydrogen band (R813, L836, and S860) efficiently abolished PigmR-mediated cell death (Figures 7F). Furthermore, either simultaneously mutating the E998-T1000-P1002 residues or deleting this predicted loop segment (residues 971-1032), also abolished PigmR-mediated cell death (Figure 7F). Therefore, these results clearly support the hypothesis that the binding ability of 2’,3’-cAMP is required for PigmR-mediated immunity.

We previously reported that another CNL R4 in the *Pigm* locus harboring 4 amino acids mutations in its LRR domain (Figure S8F), which lost rice blast resistance ^34^. To investigate whether the loss of R4-mediated blast resistance is associated with 2’,3’-cAMP binding, we performed molecular dynamics simulations of 2’,3’-cAMP interactions with R4, PigmR and other CNLs. The resulting trajectories showed small fluctuations in root mean square deviation (RMSD) for the interaction mode, indicating the high confidence in the results (Figure S8G). Binding pocket analysis further distinguished the two class of NLRs: PigmR-2’,3’-cAMP, Piz-t-2’,3’-cAMP, Pi9-2’,3’-cAMP showed larger pocket volumes (1314.366 Å, 1275.807 Å and 1771.368 Å) and higher score (0.631, 0.547 and 0.364) (Figure S8H), whereas R4-2’,3’-cAMP and Pish-2’,3’-cAMP displayed smaller pocket volumes (360.597 Å and 333.348 Å) and lower scores (0.217 and 0.103) (Figure S8I). Consistent with our binding affinity data (Figure 7A, S8B and S8J). These results suggest that PigmR and its functional paralogous NLRs possess a larger, more favorable binding pocket that enhances 2’,3’-cAMP binding affinity. Collectively, we conclude that the LRR domains of the executor CNLs conferring broad-spectrum blast resistance perceive 2’,3’-cNMP that filamentous HMA proteins generate, linking HMA-mediated second messenger production to CNL-driven immune activation.

## Discussion

HMA-containing proteins have long been identified with unrecognized molecular functions, including those HMA domain-only proteins^30,31,51^. Here we report that the NLR PigmR and its cognate effector AvrPigm define a previously unappreciated immune mechanism that depends on HMA sensor proteins with a new detection model (Figure 7H), which is likely distinct from to the typical Decoy and Guardee models ^1,52^. During infection, the conserved MAX effector AvrPigm is perceived by specific HMA proteins. Detection of AvrPigm promotes HMA proteins cytoplasmic translocation to generate 2’,3’-cNMP signaling molecules in a PigmR-dependent manner with a feedback route. These molecules are subsequently perceived by the LRR domains of PigmR, thereby initiating PigmR-mediated broad-spectrum blast resistance.

### AvrPigm-PigmR indirect interaction and durable blast resistance

Pathogens evade host perception and immune activation through rapid effector mutation ^53^. In the Rice-*M. oryzae* pathosystem, MAX effectors frequently evolve new alleles or lose function to escape recognition by NLRs ^31,42,54^. Plants have evolved cognate immunity receptors to recognize pathogen effectors and execute immunity response ^42,55^. We discover that AvrPigm exhibits multiple copies in natural populations. Moreover, single amino acid mutation did not alter its ability to trigger PigmR-mediated immunity; only complete deletion disrupted PigmR-mediated resistance. These unique features ensure the AvrPigm conservation in natural *M. oryzae* populations and the PigmR-mediated broad-spectrum and durable blast resistance ^34^. The AvrPigm-PigmR indirect interaction also defines a new model of immunity: a MAX effector is sensed by non-NLR sensors/virulence targets, and subsequently activated NLR-mediated immunity. Therefore, we provide a genetic paradigm how an Avr-NLR coevolution sustains host durable resistance in modern crops ^34^.

### HMA proteins generate 2’,3’-cNMP

We revealed the HMA proteins assemble with nucleic acids to form filament structure for catalyzing the production of 2’,3’-cNMP. Despite the architectural hints of single-strand preference, HMA proteins may also form filaments with and degrade dsDNA (Figures S3D and S4F), suggesting that these proteins might engage diverse nucleic-acid substrates in *planta.* A TIR protein, L7^TIR^, was also discovered to form a filament with dsDNA/RNA ^9^. However, the HPP04^HMA^-DNA/RNA filament is different from L7^TIR^ in the nucleic-acid components and the interaction mode: L7^TIR^ filament comprises two protein protofilaments and two dsDNA/RNA protofilaments, forming a spacious inner tunnel (40 Å diameter); in contrast, the HPP04^HMA^ filament consists of three protofilaments creating a narrower channel (14.5 Å diameter) that accommodates a ssDNA/RNA strand (Figure S4G). The shared mechanism is that filamentous structure formation is required for both proteins’ enzymatic activity.

The 2’,3’-cNMP is secondary messenger for immune response in both animals and plants ^56,57^. How 2’,3’-cNMP is generated in sufficient amount to activate immunity has been a key question in plant immunity, given that only TIR proteins were reported to deploy 2’,3’-cNMP synthetase activity, besides their NADase activity ^7,9^. Particularly in monocots which have completely lost TNLs and only retain a few TIR domain-containing proteins, and only one TIR-only protein was identified to show NADase to produce pRib-AMP/ADP ^16^. Our study reveals that the filamentous HMA proteins catalyze the production of 2’,3’-cNMP. The functionality of HMA in 2’,3’-cNMP production likely coordinates with TIR-only proteins to build the immune signal network in cereals, fine-tuning surveilling networks to avoid autoimmunity ^16,58^. Notably, the discovery that HMA proteins possess 2’,3’-cNMP synthetase activity may fill in the gap of these immune molecules’ origin and also reflects the ingenious machinery in plant immunity.

### New pair of sensor and executor proteins in plant immunity

The facts that HMAs are targeted by MAX effectors and also act as noncanonical 2’,3’-cNMP synthetase essentially extends the catalog and principle of plant immune sensor receptors, given that sensor receptors are usually NLRs that directly perceive pathogen effectors ^25,42,48,59^. Our findings highlight the role of the unique MAX effectors in ETI activation, which are also secreted by some other fungal pathogens, such as *Colletotrichum orchidophilum* and *Venturia inaequalis* ^60,61^. The ‘Arms race’ between MAX effectors and HMA-NLR module might favor the host survival with broad-spectrum resistance by deploying a repertoire of effector sensors and executor NLRs.

LRR domains of NLRs have long been expected to directly bind effectors or host guardees ^12,13,48^. We now reveal that the LRR domains of PigmR and other paralogous NLRs could perceive 2’,3’-cNMP. PigmR LRR domain forms a hydrophobic-aromatic binding pocket enriched in conserved aromatic and polar residues, which creates a specific microenvironment that selectively binds 2’,3’-cNMP to mediate immunity response. Notably, this binding pocket is highly conserved across other NLRs, including Pi9 and Piz-t. A recent study also reported that inositol binding to LRR domain of SlNRC2 is essential for SlNRC2-triggered cell death ^49^. Thus, we propose that LRR domains of NLRs might perceive diverse immune signals, shedding new light on the identification of NLR-binding molecules, given that the vast majority of the NLRs do not directly bind their cognate effectors. However, we could not exclude the possibility that these NLR LRR domains also bind other proteins or immune molecules to trigger robust immunity, in association with 2’,3’-cNMP. Nevertheless, our study provides a new concept for engineering new NLRs for broad-spectrum disease resistance in crops.

### Limitations of the study

We have identified a novel and conserved MAX effector AvrPigm and specific HMA proteins that produce 2’,3’-cNMP to activate CNL-mediated immunity. While we solved the high-resolution Cryo-EM structure of HMA bound to ssDNA/ssRNA, the technical difficulty of LRR-2’,3’-cNMP structures needs to be solved with more advanced techniques. It is also to be determined how other MAX-HMA interactions activate rice immunity, given that MAX effectors display vast sequence divergence. Further investigations would be necessarily to dissect the interconnection between TIR-generated pRib-AMP/ADP and HMA-generated 2’,3’-cNMP in cereals.

## DATA AND CODE AVAILABILITY

The raw genomic sequencing data have been deposited in the NCBI Sequence Read Archive (SRA; https://www.ncbi.nlm.nih.gov/sra) under BioProject accession number PRJNA1199142 and PRJNA1309009.

The cryo-EM map of H0PP04^HMA^-ssDNA/RNA have been deposited in the Electron Microscopy Data Bank (https://www.ebi.ac.uk/pdbe/emdb/) under the accession numbers EMD-67729. The atomic coordinate data of the HPP04^HMA^-ssDNA/RNA have been deposited in the Protein Data Bank(http://www.rcsb.org) under the accession numbers 21JC. Accession numbers are also listed in the key resources table. This paper does not report original code. Any additional information required to reanalyze the data reported in this paper is available from the lead contact upon request.

## ACKNOWLEDGMENTS

We would like to thank Xuehui Huang for help in BSA analysis. Wenjuan Cai for help in live-cell imaging; Xiaoyan Xu, Lianyan Jing, and Li Liu for helps in small molecules analysis and MST assays; Minhua Zhang and Xiaoxian Wu for assistance in cryo-EM sample preparation. We would like to express our sincere gratitude to Xiangyi Shi from Thermo Fisher Scientific Shanghai NanoPort for her valuable assistance with TEM Data collection. This work was supported by the National Natural Science Foundation of China (32088102 to Z.H.; 32425006 to Y.Z.; 32272513 and 32572799 to Z.W.; 32261143468 and U24A20405 to H.K.), the Chinese Academy of Sciences Strategic Priority Research Program (XDB1490000), the National Science and Technology Major Projects (2023ZD04070 to Y.W.). Y. Z. is a SANS Exploration Scholar.

## AUTHOR CONTRIBUTIONS

H.K., Y.D., Z.W., Y.Z. and Z.H. conceived the project. X.G., G.N., S.S., B.Y., Z.L., W.T. and P.H. performed most the experiments and analyzed the data. X.G. and Z.X. performed genomic analysis. X.G., G.N., Z.L. and X.W. developed materials and performed inoculation. B.Y., X.G., W.T., H.Z., M.C., N.P., P.H., D.T., H.K., and G.W. performed *M. oryzae* population analysis. X.G., G.N. performed gene identification and protein interaction assays. G.N., S.S., X.W. and M.Z. prepared the Cryo-EM sample and solved the structure. X.G., Z.L. and L.Z. performed cell viability and live-cell imaging assay. X.G., Y.G. and L.J. performed small molecules analysis. H.L. and J.L. assisted in field test and inoculation. X.G., G.N., Z.L., S.S., Y.D., Z.W., Y.Z. and Z.H. wrote the paper.

## EXPERIMENTAL MODEL AND SUBJECT DETAILS

### Plant materials and growth conditions

The rice *japonica* variety Nipponbare (NIPB), near-isogenic *Pigm* line (NIL-*Pigm*) and ZH11 (*Piz-t*) were used in this study. All rice lines, mutants and transgenic plants were grown in protected paddy fields at the Shanghai and Hainan Island experiment stations. For seedling experiments, plants were grown in controlled environment chambers under the conditions of 12-h day, 28°C, 80% humidity followed by 12-h night, 26°C, with 60% humidity. For rice blast pathogen spray inoculation, 14-day-old seedlings were grown at the greenhouse at 26°C, 14-h day and 10-h night.

### Pathogens

Rice blast fungus (*Magnaporthe oryzae*) isolates TH12, YN072313, Guy11, Sar-AD-3-5, YN716 and the TH12 mutant TM21, and a natural virulent isolate 5-2-1 collected from the natural blast nursery; and the transformant *pAvrPigm:AvrPigm*/TM21 were used in this study.

## METHOD DETAILS

### UV-mutagenesis and isolation of virulent mutants towards *Pigm*

To obtain virulent isolates overcoming *Pigm*-mediated resistance, continuous UV ray mutagenesis was performed as follows. Briefly, avirulent *Magnaporthe oryzae* TH12 spore suspension was exposed to UV radiation for mutagenesis using a UV crosslinker (CX-2000 UV Crosslinker, UVP) with default procedure at an irradiation intensity of 6×10^4^ µJ/cm^2^. The culture dish containing the spore suspension was placed in the center of the crosslinker and parameters were configured for UV-induced mutagenesis. After UV mutagenesis, the spore suspension was diluted to different concentrations and spread on complete medium agar plates to check survival rates, and the mutagenized spore suspensions were also sprayed onto NIL-*Pigm* rice seedlings in a well-protected incubator for screening virulent mutants. Initial virulent isolates were then detached from the infection lesions, and a second round of UV mutagenesis was applied to obtain complete virulent isolates. Through this mutagenesis process, 12 independent virulent mutants (designated TM) were isolated, including TM21. For natural virulent isolates, blast lesions were collected from different natural blast nurseries, and spores were collected after detaching isolates from these lesions, and were cultures and inoculated onto *Pigm* seedlings, and a natural virulent mutant, 5-2-1, was finally identified from hundreds of natural lesions.

### DEB (Diepoxybutane)-mutagenesis and isolation of virulent mutants towards *Pigm*

Step 1. Protoplasts treatment with Diepoxybutane (DEB)

The NIL-*Pigm* avirulent strain YN716 was cultured on oatmeal agar (OAT) medium under continuous light to induce conidiation. Conidia were harvested and inoculated into liquid complete medium (CM) for the formation of mycelial balls. Protoplasts were then released from these mycelial balls via enzymatic digestion, using a solution of 0.075 g Megalyase dissolved in 10 ml of 0.7 M NaCl. The protoplasts were washed and resuspended in 0.7 M NaCl for subsequent use.

Mutagenesis was conducted by adding the mutagen diepoxybutane (DEB) to the protoplast suspension, adjusting to a final concentration of 0.001%. The mixture was gently inverted to ensure uniform mixing and then incubated in the dark for 36 hours. Post-mutagenesis, the protoplasts were washed with 0.7 M NaCl to eliminate residual DEB, and resuspended in 5 ml of liquid TB3 medium for recovery. The recovery culture was incubated at 28°C with shaking at 80 rpm for 12∼16 hours.

Regenerated cells were mixed with 45°C unsolidified OAT medium to a final volume of 50 ml, and the mixture was poured into 12 cm Petri dishes pre-filled with a bottom layer of plain oatmeal agar. After the medium solidified, the plates were first incubated in the dark for 2 days, followed by 7 days of incubation under continuous light to facilitate sporulation.

Step 2. Isolation of virulent strains towards *Pigm*.

Conidia from the resulting mutant colonies were collected and sprayed onto three-leaf-stage NIL-*Pigm* rice seedlings. Seven days post-inoculation, leaf lesions were sampled, and single conidia were isolated from these lesions using a microscope. The isolated strains were cultured, and their pathogenicity was verified in accordance with Koch’s postulates.

### *AvrPigm* cloning integrating map-based cloning, bulked segregant analysis and PacBio RSII re-sequencing

A genetic mapping population was constructed by sexual crossing the virulent isolate TM21 with the avirulent isolate YN8773R-27, yielding 111 progenies ^62^. Next, each progeny was then individually cultured and inoculated onto the susceptible cultivar NIPB or the resistant cultivar NIL-*Pigm* to assess virulence phenotypes.

For map-based cloning and Bulked Segregant Analysis (BSA), genomic DNAs of *M.oryzae* were extracted using the cetyl trimethyl ammonium bromide (CTAB) method. Genomic DNA libraries (prepared from parental isolates and F_1_ progenies were sequenced for paired-end (PE) short reads on the Illumina HiSeq platform (Illumina, California, USA). Raw sequencing reads were aligned to the TH12 reference genome using BWA v0.7.17 with default parameters. Genetic variants were obtained using BCFtools v1.10.2 and manhattan plot was generated with qqman.

For long-read sequencing, the extracted DNAs were purified with Grandomics Genomic DNA Purification Kit following the manufacturer’s standard protocol. Sequencing was performed by Grandomics (Wuhan, China) on the PacBio Revio platform. Collinearity analysis was visualized by gggenomes.

To identify sequences related to AvrPigm, tblastn searches were performed against both published ^40,41^ and newly re-sequenced *Magnaporthe oryzae* genomic datasets, using default parameters for all analyses. Pogenies inoculation and *M.oryzae* genomic data are listed in Table S1.

### *M. oryzae* genetic transformation

To perform genetic complementation of *AvrPigm* in mutant TM21, the coding sequence of *AvrPigm* along with its upstream 2 kb promoter region and downstream 0.5 kb region was cloned into the modified vector pCambia 1300. *M. oryzae* transformation was performed as described previously ^58^. Briefly, a single colony of *Agrobacterium tumefaciens* AGL1 harboring the *AvrPigm* expression construct was picked into 5 ml liquid LB medium and shake culture at 28°C, 200 rpm for 2 days. Next, 1 ml of AGL1 bacterial culture was transferred into 10 ml IMAS (Induction Medium Acetosyringone) liquid medium and shake culture at 28°C, 200 rpm for 6 hours. Bacterial suspension (OD_600_, 0.5) using IMAS liquid medium. IMAS liquid medium washed *M. oryzae* spores and adjust the concentration to 1 × 10^6^ spores/ml. Mix equal volumes of *Agrobacterium* liquid and spores. Pipette 200 µl of the mixture onto sterile filter paper placed on solid IMAS medium and co-cultured under light-avoiding conditions for 3 days. The filter paper was then transferred into complete medium (CM) supplemented with hygromycin, cefotaxime sodium, and carbenicillin-to select for transformants. After 3 days, DNA of transformants was extracted to identify the positive transformants. More than 15 independent isolates were selected and their spores were sprayed onto NIPB (susceptibility) and NIL-*Pigm* (resistance).

For subcellular localization analysis of AvrPigm and Avr-Piz-t, the protoplast-mediated transformation of *M. oryzae* was performed ^63^, and the AvrPigm:GFP and the Avr-Piz-t:mCherry constructs were introduced into the protoplasts of wild type Guy11. The resulting transformants were screened by using fluorescence microscopy. The fluorescent strains (*pAvrPigm*:AvrPigm:GFP and *pAvrPiz-t*:AvrPiz-t:mCherry) were selected to perform plant infection. For inoculation, spores were harvested and resuspended to a final concentration of 1 ×10^5^ spores per milliliter. A spore suspension was injected into rice sheaths (NIPB and NIL-*Pigm*) followed by incubation at 28°C for 30 h under high humidity (90%). Epidermic cells of rice sheaths were observed and photographed under fluorescence microscopy. The confocal microscopy (Nikon TiE system Nikon, Japan) was used for fluorescence imaging. Primer sequences are available in Table S4

### Pathogen inoculation experiments

Blast fungal inoculation was conducted as previously described ^36^. Briefly, *M. oryzae* isolates were cultured on CM solid medium at 25°C for about 10 days to generate spores. Newly expanded leaves at tillering stage were punch-inoculated with spores diluted in water at a concentration of 1 × 10^5^ spores/ml. For spray inoculation of seedlings, two-week-old seedlings were sprayed with rice blast spore suspensions. Inoculated seedlings were kept in an incubator at 26°C for 24 h in darkness, followed by 26°C with 12 h/12 h (day/night) and 90% relative humidity for 7 days. Lesion lengths or areas were measured by ImageJ and fungal growth was determined using quantitative PCR, employing OsActin as the reference genes for rice leaves and 28s rDNA for *M. oryzae*.

### Development of rice transgenic materials

For HMAs (HPP04, HIPP16, HIPP19 and HPP04) overexpression, the coding regions of HMAs fused with FLAG tag were cloned into the vector pUN1301, and were transformed into NIPB, NIL-*Pigm* and ZH11 separately. For the inducible expression of estradiol-induced *HPP04*, the coding region of HPP04 fused with FLAG tag was inserted into the vector pBWAHVE, and subsequently transformed into NIPB and NIL-*Pigm* respectively. For compartmentalized expression of HPP04, signal peptides (MYR, NLS, NES) were fused to the N-terminal of HPP04, and a FLAG tag was fused to the C-terminal. The resulting constructs were cloned into the vector pUN1301 and transformed into NIPB and NIL-*Pigm*, respectively.

For CRISPR/Cas9 knockout construct, aimed target sequences of HIPP16, HIPP19, HIPP20, HPP04 were chemically synthesized and ligated into pOs-sgRNA, which were inserted into the transformation vector. For the multiple knockout constructs, the sgRNAs were chosen according to the manual and inserted into the transformation vector. All primer sequences used for vector construction are listed in Table S4.

### Phylogenetic tree and yeast two hybrid (Y2H) assay

The HMAs family proteins were identified by HMMER’s hmmsearch tool based on Hidden Markov Model (HMM) from Interpro (Pfam) based NIPB genomes (RAP-DB) and already published data ^29^. The maximum likelihood tree was constructed by MEGA X with 1,000 bootstrap replicates. The functional domain annotation was carried out by hmmscan, designed the database was pfamA, cut-off evalue is 1e^-5^. Rice HMA protein family members are listed in Table S3.

To identify AvrPigm-interacting HMA domain-containing proteins (HMAs), signal peptide-truncated cDNA fragment of AvrPigm was inserted into pGBKT7 (bait) vector (Clontech, http://www.clontech.com/). Concurrently, full-length coding sequences of HMAs family proteins were inserted into pGBKT7 (prey) vector. Paired construct group were co-transformed into yeast strain AH109 via polyethylene glycol/lithium acetate (PEG-LiAc) method. Transformed yeast cells were then plated onto synthetic dropout (SD) medium lacking Trp, Leu, His, and adenine and containing proper 3-aminotriazole to select for positive transformants.

### Recombinant protein expression and purification

Full-length coding sequence of HMAs, HMA-containing proteins and AvrPigm without signal peptide were cloned into pET28a-6 × His-SUMO2 vector. All the constructs were expressed in *E. coli* BL21 (DE3). The cell cultures were grown at 37°C to OD_600_ of 0.6-0.8. Then the cultures were cold shocked on ice for 30 min before adding Isopropyl-β-D-thiogalactoside (IPTG) to induce protein expression at 16°C for 16 h. The *E.coli* pellets were collected and lysed in lysis buffer containing 25 mM Tris-HCl at pH 8.0, 500 mM NaCl, 5% glycerol, 50 mM Glycine, 2 mM MgCl_2_, 5 mM β-mercaptoethanol. The lysis mixture was centrifuged at 15,000 g for 1h. The supernatant was loaded onto a Ni-NTA affinity column (Cytiva) equilibrated with lysis buffer. After washing the column three times with 20 mM imidazole in lysis buffer, SUMO2 fusion proteins were cleaved by supplementing elution fractions with approximately 250 µg of human SENP1 protease during 3 hours at 4°C in lysis buffer with resin, then eluting with 300mM imidazole in lysis buffer to obtain no-tagged proteins. The purified proteins were dialyzed overnight against a storage buffer (20 mM Tris-HCl at pH 8.0, 100 mM NaCl, 50 mM Glycine, 2 mM MgCl_2_). Further purification was carried by size-exclusion chromatography using Superose6 10/300 gel filtration column in dialysis buffer.

The LRR domains of NLR (PigmR, Pi9, Piz-t, and Pish) were inserted into pCold-TF (TaKaRa) with N terminal 6 × His and transformed into *E.coli* BL21(DE3). After obtaining the eluted protein solution, the proteins were dialyzed overnight against a buffer containing 20 mM Tris-HCl pH 8.0, 100 mM NaCl, 1% glycerol, 5 mM β-mercaptoethanol. Through Ultrafiltration obtain the aimed proteins.

### Negative-stain electron microscopy

For quality examination, samples for negative staining were prepared on 400 copper mesh grids that were glow-discharged with negative polarity, 15 mA for 60 s, using a PELCO easi-Glow glow discharge system. HPP04^HMA^ (6 µl) was deposited on the grid. After incubation of 60 s, excess sample was removed from the grid by touching the edge of a filter paper. After the blotting, the grid was floated 3 times in 2% uranyl formate solution for 5 s, then dried at room temperature for 2 min. Negative-staining micrographs were recorded using a Hitachi H-7700 120 kV transmission electron microscope (TEM) and images were recorded with an AMT XR-41 camera.

### Cryo-EM sample preparation and data collection

The HPP04^HMA^ protein (3 µl, 0.5 mg/ml) was applied to Quantifoil R1.2/1.3 300 mesh girds, which were glow-discharged at 15 mA current and 0.26 mBar vacuum pressure for 60 s using a glow-discharge cleaning system (PELCO easiGlow). The grids were vitrified in a FEI Vitrobot Mark IV (Thermo Fisher Scientific) using a blotting time of 1 s, blot force -1, 8 °C, and 100% humidity, with a double-layered TED PELLA filter paper was used during blotting.

Cryo-EM data were collected on a Krios G4 electron microscope (equipped with a Falcon 4 Direct Electron Detector, a Selectris X Imaging Filter with a slit width 10 eV; Thermo Fisher) in counting mode with Compression on, which was operated at ×165,000 magnification with a physical pixel size of 0.37 Å. EPU software was used for data collection, with an exposure rate of 6.46 electrons per pixel per second. The micrographs were dose-fractionated into 40 frames, resulting in a total exposure time of 4.66 seconds and a total electron exposure of ∼ 55 electrons per Å^2^. The defocus values range from - 1.4 to -1.6 μm.

### Cryo-EM data processing and model building

A total of 14,222 movies were 2×Fourier-binned and motion-corrected using MotionCor2 ^64^ within RELION 3.1 ^65^, resulting in a pixel size of 0.74 Å per pixel. The defocus values of the motion-corrected micrographs were then estimated using Patch CTF in cryoSPARC v4.2.1 ^66^. Unless otherwise stated, all subsequent processing steps were performed using cryoSPARC v4.2.1. From the motion-corrected micrographs, 214 particles were manually picked from 68 randomly selected images to generate several templates. A total of 1,603,803 particles were then picked using Filament Tracer (filament diameter = 60 Å) and extracted with a box size of 256 pixels. These particles underwent two rounds of 2D classification (N=100, Iterations=35, align filament classes vertically, do not remove duplicate particles). An initial model was generated using Ab Initio Reconstruction, followed by Homogeneous Refinement without imposing symmetry. The particles were then subjected to 3D classification (N=10). The six 3D classes exhibiting similar conformational features were selected and merged. These particles were further refined by homogeneous refinement followed by NU refinement. The final reconstruction displayed clear structural features of the HPP04^HMA^ protein at a resolution of 3.54 Å.

The predicted structure of HPP04^HMA^ by AlphaFold3 ^50^ was fitted to the EM density in Chimera to build the initial model. Local variations were manually adjusted in Coot v0.9.8.7 based on the global map with torsion, planar peptide, trans peptide, and Ramachandran restrains ^67^. Multiple rounds of real-space refinement in Phenix were used to refine the model with secondary structure and Ramachandran restrains (weight =0.25) ^68^. Details of the model statistics can be found in table S3.

### Transient gene expression and subcellular localization assay in *N.benthamiana*

The tested full length or domain sequences were cloned into pCAMBIA-35S vectors with different tags, including Myc and GFP. Plasmids were transformed into *Agrobacterium* strain GV3101, cultured overnight in LB medium, collected and resuspended in infiltration buffer (10 mM MgCl_2_, 10 mM methylester sulfonate, 150 µM acetosyringone, pH 5.6), and incubated for 2-3 h at 30°C before infiltration. Adjusting the concentration of GV3101 to OD_600_ = 0.8. The suspensions were then infiltrated into 5-week-old *N.benthamiana* leaves in various combinations. Each group covers approximately 1-2 square centimeters. After 36–48 hours, the infiltrated leaves were collected and the protein expression was identified via western blotting. Western blotting was imaged using a Tanon-5200 imaging system (Tanon).

### Protoplasts transformation and cell viability

Constructs of HMAs and AvrPigm were cloned to pA7-35s-Flag-RBS, pA7-35s-HA-RBS or pUC19-35s-eGFP-RBS with additional Xlinker for protoplasts transfection. Rice protoplasts were prepared from leaf sheaths of 12-day-old NIPB or NIL-*Pigm* seedlings. Protoplasts were transfected with the constructs via polyethylene glycol (PEG)-mediated transformation for 12 h. Protoplasts were collected and rapidly frozen in liquid nitrogen, and stored at -80°C for subsequent protein extraction, or used immediately for downstream assays.

To assess protoplasts viability following transfection, the different group transfected with indicated plasmids were incubated for 12 h. Cell viability was determined by the Cell Titer-Glo luminescent Cell Viability according to the manufacturer’s instructions (meiluncell, PWL111). Luminescence intensity - proportional to intracellular ATP levels was measured by Varioskan Flash multireader (Thermo Fisher Scientific). The tested cells were normalized against the control, assigned as 100%, to calculate the survival percentages. Proteins were extracted via UREA-based method. Briefly, protoplasts were harvested by centrifugation at 200 ×g for 5 min. The pellet was washed twice with WI solution to remove excess calcium ions. Discarding the supernatant, the pellet was suspended in 100 µl of extraction buffer (5% SDS, 4 M urea, 0.2% Triton X-100, 100 mM DTT, pH 8.0) and vortexed for 5 minutes. The suspension was then frozen in liquid nitrogen for 1 minute and thawed in a 37°C water bath until fully thawed. This freeze–thaw cycle was repeated three times to enhance protein release. After the final thaw, 5×SDS loading buffer was added to the samples. The samples were then incubated at 95°C for 5 minutes, followed by centrifugation at 12,000 rpm for 10 minutes. The supernatant is collected for subsequent SDS-PAGE.

### Confocal microscopy for protein localization

For *Agrobacterium*-mediated transient expression in *Nb*, leaves were imaged for protein localization analyses between 36 h and 48 h post infiltration. Confocal images were taken on a confocal laser scanning microscope SP8 from Leica (Germany). GFP was excited using a 488-nm laser, and the emission spectrum was detected between 516 and 556 nm; mCherry was excited with a 532-nm laser, and the emission spectrum was detected between 570 and 620 nm. The number of puncta per 50 um PM was counted via ImageJ. Statistical analysis was performed with student’s t.test or Turkey’s HSD test, with asterisks and different letters indicating significant differences.

For observation of subcellular localization of aimed proteins in rice protoplasts, the indicated constructs were transformed and incubated for 12 h. Imaging was performed using the same Leica SP8 confocal microscope and laser settings described above to ensure consistency in fluorescence detection and analysis.

### Pull-down and co-immunoprecipitation assay

For pull-down assay, recombinant proteins were generated using a dual-expression vector system. Briefly, the His-tagged HMAs and AvrPigm signal peptide-truncated form tagged with S-tag were inserted into two separate multi cloning sites in the vector pet Duet. The supernatant containing soluble proteins were collected and purified by Ni-NTA resin. After washing, the eluted proteins were separated and analyzed by SDS-PAGE. Checking the purpose proteins by Coomassie brilliant blue and western blot.

For co-immunoprecipitation assay, the indicated constructs were under control of the 35S promoter and transformed into rice protoplasts. After incubating for 12 h, and the total proteins were extracted with Co-IP buffer (25 mM Tris-HCl at pH 8.0, 100 mM NaCl, 5% glycerol, 0.5 % Triton X-100, protease inhibitor cocktail). After centrifugation for 30 min, the supernatants were incubated with anti-Flag magnetic beads for 2 h at 4 °C and then washed three times with Co-IP buffer. The bound proteins were eluted from the beads using a flag peptide competition and analyzed by western blotting.

### Electrophoretic mobility shift assay

Cy5-labeled DNA probes (5’-end labeled) were chemically synthesized to detect interactions with the HPP04 protein. The purified HPP04 protein (25-2000 nM) was incubated with the Cy5-probes at 4 °C for 8 h in EMSA buffer (20 mM Tris-HCl pH 8.0, 100 mM NaCl). Following incubation, the reaction mixtures were stopped and electrophoresed in 4.5% native polyacrylamide gel. Labelled probes in the gel were visualized using a Typhoon scanner (Fujifilm) to capture fluorescence signals.

### In vitro nuclease activity assay

Rice genomic DNA and total RNA samples (200 ng for each) were incubated with purified HIPP16^HMA^, HIPP19^HMA^, HIPP20^HMA^, HPP04^HMA^ and RGA5^HMA^ (2 µM for each) proteins separately in the reaction buffer including 25 mM Tris-HCl pH 8.0 and 150 mM NaCl at 25 °C. After reaction for 16 h (for DNA), the reaction mixtures were mixed with DNA loading buffer and visualized by agarose gel electrophoresis. For RNA degradation, samples were collected 6 h after incubation and visualized by agarose gel electrophoresis in a similar way.

Nuclease activity was further measured by DNase І Assay Kit (Fluorometric, Abcam) with a commercially fluorescent DNA probe as the substrate. The purified proteins (5 µM) individually were incubated with the fluorescent DNA probe (25 µM, Abcam) in the assay buffer. The samples were measured by Varioskan Flash multireader (Thermo Fisher Scientific) with excitation/emission 651/681 nm fluorescence.

### High-performance liquid chromatography-mass spectrometry analysis

Samples prepared as described above were analyzed Q Exactive quadrupole orbitrap high resolution mass spectrometry coupled with a Dionex Ultimate 3000 RSLC (HPG) ultra-performance liquid chromatography (UPLC-Q-Orbitrap-HRMS) system (Thermo Fisher Scientific), with a HESI ionization source. Samples were separated with an ACQUITY UPLC HSS T3 column (100×2.1 mm, 1.8-μm particle size; Waters). The mobile phase consisted of (A) water supplemented with 2 mM ammonium acetate and (B) acetonitrile. The gradient of B was as follows: 0.0-2.0 min, 2.0%; 2.0-6.0 min, 2.0% to 60%; 7.0-8.0 min, 99.0%; 8.1-12.0 min, 2%. The flow rate was 0.30 mL/min and column temperature 40 °C.

All MS experiments were performed in negative ion modes using a heated ESI source. The source and ion transfer parameters applied were as followed: spray voltage 3.0 kV(negative). For the ionization mode, the sheath gas, aux gas, capillary temperature and heater temperature were maintained at 40 psi, 10 psi, 320 °C and 350 °C, respectively. The Orbitrap mass analyzer was operated at a resolving power of 70,000 in full-scan mode (scan range: 100-1050 m/z; automatic gain control (AGC) target: 1e^6^) and of 17,500 in the Top 5 data-dependent MS^2^ mode (stepped normalized collision energy: 20, 40 and 60; injection time: 50 ms; isolation window: 1.5 m/z; AGC target: 1e^5^) with a dynamic exclusion setting of 4.0 seconds.

### Production and detection of 2′,3′-cNMP in vitro

Purified HIPP16^HMA^, HIPP19^HMA^, HIPP20^HMA^, HPP04^HMA^, RGA5^HMA^ and L7^TIR^ (2 µM for each) were individually incubated with 500 ng of rice genomic DNA or total RNA in reaction buffer. As controls, 2 µl of RNase T1 (Thermo Scientific) or DNase І (NEB) were incubated with the same amount of DNA or RNA with the same conditions. Following incubation at 25°C for 16 h (DNA) or 6 h (RNA), adding same volume methanol and centrifuging at 12,000 g for 10 min to remove proteins, collecting the supernatant and analyzing with LC-MS/MS for metabolite identification and quantification.

The compounds were analyzed by a QTRAP mass spectrometer (5500, AB Sciex) equipped with UHPLC (ExionLC AD, AB Sciex) and an XSelect HSS T3 column (100mm × 3.0 mm, 2.5 μm, Waters). The injection volume was 1 μl. The mobile phase A was 2 mM ammonium formate in water and B was methanol. The column was maintained at 40 °C with a flow rate of 0.4 mL min^-1^, and the gradient of B was as follows: 0 min, 2%; 1min, 2%; 8 min, 15%; 9 min, 95%; 11 min, 95%; 11.1 min, 2%; 15 min, 2%. All analytes were detected using Multiple reaction monitoring (MRM) mode, The optimized ESI operating parameters for negative mode were: ion spray voltage, 4.5 kV; ion spray temperature, 500°C; curtain gas, 35 psi; ion source gas 1, 50 psi; ion source gas 2, 50 psi. Metabolites quantification was based on peak area, and the retention time for each compound was determined using standards dissolved in sterile water. MRM conditions were optimized using authentic standard chemicals including: 2′,3′-cAMP ([M-H]- 328.00>134.00, 328.00>107.00), 2′,3′-cGMP ([M-H]-344.00>150.00, 344.00 >133.00), 2′,3′-cCMP ([M-H]- 304.00>110.00, 304.00>67.00), 3′,5′-cAMP ([M-H]- 328.10>134.00, 328.10>107.00), 3′,5′-cGMP ([M-H]- 344.10>150.00, 344.10 >133.00), 3′,5′-cCMP ([M-H]- 304.10>110.00, 304.10>67.00). Instrument control and data acquisition were performed using Analyst 1.6.3 software (AB SCIEX), and data processing was performed using MultiQuant 3.0.2 software (AB SCIEX).

### Detection of 2′,3′-cNMP in planta

2′,3′-cNMP were extracted from rice tissues follow the reported procedure^9^. In brief, rice transgenic or wild type plants growing in the green house. Corresponding stage leaves were harvested and liquid nitrogen-frozen for measurement. Frozen leaf tissues were homogenized with a tissue lyser, approximately 100 mg ground tissues were added extraction buffer containing 600 µl 4% acetic acid with 20 µl 1 mM phosphodiesterase inhibitor 3-isobutyl-1-methylxanthine (IBMX, Sigma). For protoplasts from NIPB or NIL-*Pigm*, about 1 × 10^6^ rice protoplasts were collected and resuspended in the same 600 µl extraction buffer. The samples (tissues or protoplasts) were vortexed for 2 min and subsequently centrifuged for 10 min at 12,000g. The supernatants were transferred into new pre-cooled 1.5 ml tubes. 800 µl chloroform was added to each tube, and vortex equally for 2 min and centrifuged at 6,000g for 10 min. Aqueous phases were collected and extracted again by chloroform. The samples were dried under vacuum concentrator to enrich the small molecules. Dried pellets were reconstituted in 50 µl of water and ultrasonic resolution. The samples were centrifuged 12,000 rpm for 10 min, and the supernatants were applied to the LC-MS/MS for metabolite identification and quantification.

### Protoplast-based cell death analyses by spinning-disk confocal microscopy

The spinning-disk confocal microscopy system was used on an Olympus IX83 microscope equipped with a X-Line Extended Apochromat Infinity Corrected 40× Objective (0.95NA, Olympus) and OBIS 488 nm LS 150 mW, 561 nm LS 150 mW and 640 nm LX 150 mW lasers (Coherent), a CSU-W1 spinning disk confocal unit (Yokogawa), and a CCD camera (ORCA-Fusion BT, Hamamatsu). Imaging sequences were acquired by CellSens Dimension (Evident).To track the dynamic process of AvrPigm and HMA proteins triggered cell death in NIL-*Pigm*, protoplasts were transfected with different plasmids for 6 h and then incubated with 100 ng/ml of SYTOX^TM^ Blue Dead Cell Stain (Thermo Fisher Scientific). The protoplasts in culture medium were then moved to a 20 mm glass bottom dish (NEST), filled with 2 ml medium and imaged by the spinning disk confocal microscopy. The auto fluorescence in protoplasts was excited by the 640-nm laser, and the dye signal was excited by the 405-nm laser, which is also used to indicate the morphology of the whole protoplasts in bright field. The protoplasts were imaged every 30s (exposure time 200 ms) from 30 min to several hours for time-lapse videos. The generation of montages, measurement of cell areas, measurement of fluorescence intensities, applying the LUT of images, and merge of channels were done by Fiji.

### Subcellular fractionation isolation in rice

Subcellular fractionation was performed using the Minute™ Plasma Membrane Protein Isolation Kit (Invent; Cat# SM-005-P) following the manufacturer’s protocol. Following fractionation, isolated fractions were resuspended in LDS loading buffer (Tanon; Cat# 180-8210D) and denatured by heating at 95°C (70°C for PM fractions) for 5 min. Subsequently, total, nuclear extract, cytosolic fraction, and plasma membrane fraction were resolved on SDS-PAGE. Proteins were then transferred to PVDF membranes for immunoblotting with specific antibodies.

### Microscale thermophoresis (MST) analysis and Flow-induced dispersion analysis (FIDA)

Binding affinities of recombinant His-TF-R4^LRR^, His-TF-PigmR^LRR^, His-TF-Piz-t^LRR^, His-TF-Pi9^LRR^, and His-TF-Pish^LRR^ to 2′,3′-cAMP/cGMP were measured by microscale thermophoresis in a Monolith NT.Label-Free instrument (Nano Temper Technologies GMBH, Germany). We also measure the binding affinities between HMA proteins and AvrPigm/ssDNA/ssRNA. The above proteins (20 nM) were prepared in assay buffer (1 × PBS buffer containing 0.05% (v/v) Tween-20). A range of concentrations of 2′,3′-cAMP/cGMP in the assay buffer were incubated with labeled protein (1:1, v/v) for 5 min. The sample was loaded into the NT.Label-Free standard capillaries and measured with auto-detected LED power and MST power. The KD Fit function of the Nano Temper Analysis Software was used to fit the curve and calculate the value of the dissociation constant (*K*d).

Binding studies of 2′,3′-cAMP/cGMP to purified LRR domains of NLRs were performed using Flow induced dispersion analysis (FIDA) on a Fida 1 instrument equipped with LED-induced fluorescence detection with 280 nm detector (Fida Biosystems ApS, Copenhagen, Denmark). All experiments utilized coated capillaries (Cat100-002). Proteins were maintained at a fixed concentration of 0.5 μM, while the analyte (2′,3′-cAMP) was titrated from 50 nM to 1 mM in the 1× PBS assay buffer. The indicator-analyte capmix method was performed in the capillary during measurement at 400 mbar run pressure for 180 s. The sample tray was incubated at 4°C throughout the experiment and all samples were measured in technical triplicates at 25°C. Data was analyzed using FIDA software version V2.36 (Fida Biosystems ApS, Copenhagen, Denmark) to extract binding parameters.

### Native MS analysis to characterize 2′,3′-cNMP interactions with LRR domains of NLRs

To determine the native state binding of 2′,3′-cNMP to the LRR domains of NLR proteins, samples were prepared under non-denaturing conditions to preserve protein folding, adapted from (Ma et al. 2024). The LRR domains of PigmR, Pi9, Piz-t, and Pish were adjusted to a final concentration of ∼4 μM in the buffer containing 10 mM Tris-HCl (pH 8.0), 100 mM NaCl with 40 µM 2′,3′-cNMP. Samples were incubated overnight at 4°C to allow equilibrium binding of 2′,3′-cNMP to the LRR domains. Next, anti-His affinity resin was added to the protein-2′,3′-cNMP mixtures and incubate for 2 h at 4°C to capture the complexes. The resin was pelleted by centrifugation at 3,000 ×g for 10 minutes at 4°C, and the supernatants (containing unbound 2′,3′-cNMP and contaminants) were discarded. The resin was washed three times with 1 ml of wash buffer (10 mM Tris-HCl, pH 8.0, 100 mM NaCl) to remove non-specifically bound molecules. The LRR-2′,3′-cNMP complexes were eluted using 300 mM imidazole (pH 8.0) to disrupt His-tag-resin interactions. The samples were added with same volume methanol to precipitate proteins and centrifuged at 15,000 ×g for 30 min. The supernatants were dried under vacuum concentrator and resuspended in sterile water. The final supernatants were applied to the LC-MS for metabolite identification and quantification.

### Structure prediction and Molecular dynamics simulation

The 2′,3′-cAMP–PigmR and other NLRs complex was constructed by AF3 with default parameters. And the binding residues were analyzed by LigPlot.

The molecular dynamics (MD) simulations were performed using GROMACS 2022.1 with the Amber14sb force field under periodic boundary conditions in an isothermal-isobaric ensemble. We used the docking pose as initial structure of the complex. The molecular system was solvated in a cubic box of 8.0 ×8.0 ×8.0 nm³using the TIP3P water model to hydrate the protein-ligand complex. All atoms were parameterized with the Amber14sb all-atom force field.

The initial structure underwent energy minimization through 50,000 steps of steepest descent algorithm to eliminate steric clashes. Subsequent simulations employed the NPT ensemble with equations of motion integrated using the Leap-Frog algorithm. Long-range electrostatic interactions were calculated via the Particle Mesh Ewald (PME) method, while van der Waals and Coulombic interactions were truncated at 12 Å with neighbor list updates every 10 steps. Bond lengths were constrained using the LINCS algorithm with parameters lincs_iter = 1 and lincs_order = 4.

The system temperature was gradually increased from 0 K to 298.15 K using the V-rescale thermostat, with pressure maintained at 1 bar through the Parrinello-Rahman barostat under isotropic conditions. A dual-range neighbor searching scheme was implemented with short- and long-range cutoff distances of 9 Å and 14 Å, respectively. Hydrogen bond formation criteria required donor-acceptor distances < 0.35 nm and donor-hydrogen-acceptor angles < 30°.

Initial velocities were assigned according to Maxwell-Boltzmann distribution. The production run consisted of 50,000,000 steps with a 2 fs time step, yielding 100 ns of total simulation time and 10,000 trajectory frames. Trajectory analysis and visualization were performed using native GROMACS utilities and VMD software. All simulations maintained energy conservation errors below 0.005 kJ·mol⁻¹·ps⁻¹, ensuring numerical stability throughout the 100 ns sampling period. Technical support provided by Phadcalc.

