## supplemental figures for "HMA proteins produce 2′,3′-cNMP signaling molecules and activate CNL-mediated immunity in rice": AvrPigm-Figures-supplement.pdf

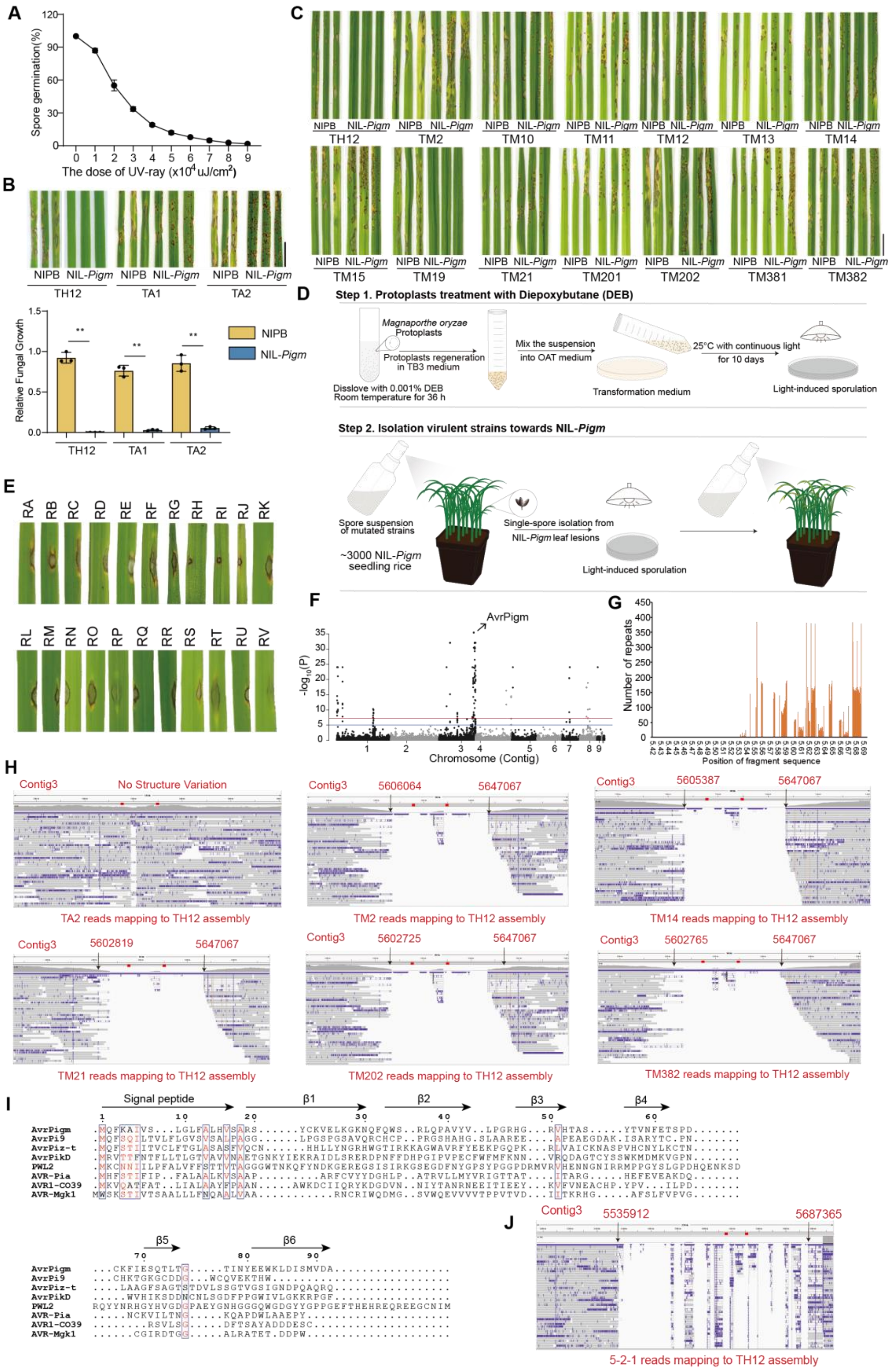

**Figure S1. Fine-mapping and characterization of *AvrPigm*, related to Figure 1.**

(A) Optimization of ultraviolet irradiation (UV) mutagenesis for *M. oryzae* spores. Maximal recovery of virulent mutants occurred at a UV dose inducing ~90% spore mortality, highlighting a narrow window for effective mutagenesis while preserving fungal viability.

(B) A single round of UV mutagenesis resulted in only partially virulent isolates and failed to generated fully virulent strains towards *Pigm*. Scale bar, 1 cm.

(C) 12 independent virulent isolates generated through two consecutive rounds of mutagenesis caused typical disease symptoms on NIL-*Pigm*. TH12 and TM19 were avirulent isolates towards *Pigm*. Scale bar, 1 cm.

(D) DEB-mediated mutagenesis of *M. oryzae* also yielded isolates virulent to *Pigm*, with a schematic illustrating the mutagenesis procedure.

(E) 22 independent isolates generated by DEB mutagenesis were virulent to *Pigm* and caused clear disease symptoms.

(F) Bulked segregant analysis (BSA) of *AvrPigm* using an F1 progeny population derived from a genetic cross between Genetic association of the TM21 and YN8773R-27 identified a major association peak on telomere of Contig (chromosome)3, corresponding to *AvrPigm*.

(G) Repetitive sequences impede fine-mapping of *AvrPigm*. The candidate *AvrPigm* interval contains abundant repetitive DNA elements.

(H) Structural variation in the telomeric region of chromosome 3 harboring *AvrPigm*. Integrative Genomics Viewer (IGV) revealed the similar large deletions overlapping the *AvrPigm* locus region in five independent virulent strains. The TH12 genomic sequence of TH12 was used as the reference genome. TA2, a first round UV-induced mutant, from failed mutagenesis, serves as a negative control. The red rectangle indicates the location of *AvrPigm* locus.

(I) AvrPigm lacks sequence homology to known MAX effectors. The amino acid sequences are aligned by MUSCLE.

(J) IGV revealed a large deletion spanning the *AvrPigm* locus in the natural virulent isolate 5-2-1, compared with the avirulent isolate TH12, resulting in complete loss of *AvrPigm*.

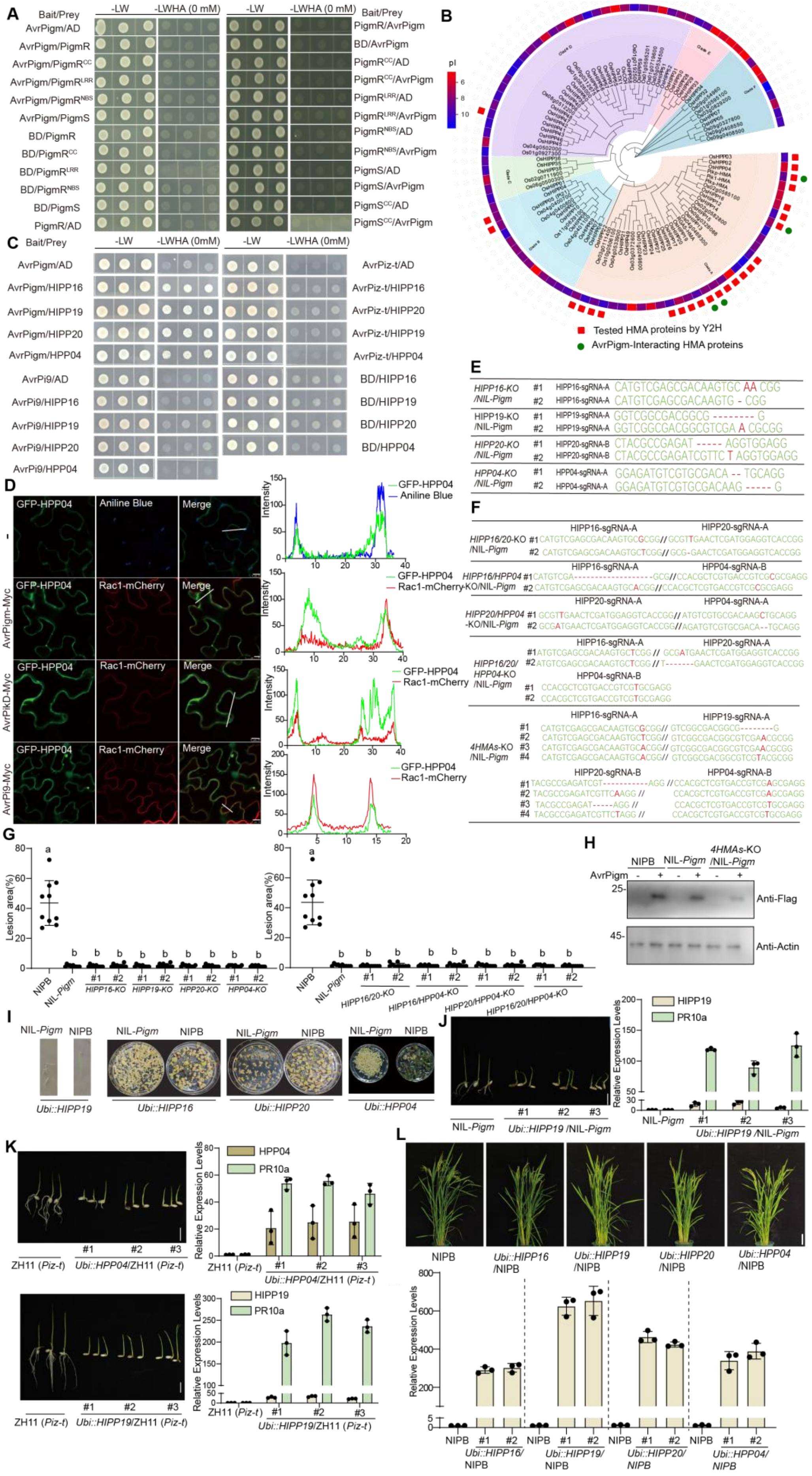
A
Bait/Prey
-LW
-LWHA (0 mM)
B
Tested HMA proteins by Y2H
AvrPigm-Interacting HMA proteins
C
D
GFP-HPP04
Aniline Blue
Merge
Rac1-mCherry
AvrPigm-Myc
AvrPikD-Myc
AvrPi9-Myc
E
F
G
Lesion area(%)
H
Anti-Flag
Anti-Actin
I
J
K
L
Relative Expression Levels

**Figure S2. Four HMA proteins associate with AvrPigm, but not PigmR, and act redundantly to activate autoimmunity in plants harboring *Pigm* or *Pizt*, related to Figure 2.**

(A) Y2H assays showed no detectable interaction between AvrPigm and full-length PigmR or PigmS, or their CC, NBS, and LRR domains. Growth on SD/–Leu/–Trp and selective SD/–Leu/–Trp/–His/–Ade medium.

(B) Phylogenetic analysis of rice HMA family proteins. A maximum-likelihood phylogenetic tree of rice HMA family proteins was constructed using MEGA-X (1,000 bootstraps). Red squares denote HMA proteins tested for interaction with AvrPigm; green circles highlight HMAs confirmed to interact with AvrPigm.

(C) Y2H assays revealed that specific interactions between four HMA proteins and AvrPigm, but not AvrPi9 and AvrPiz-t. Negative controls (empty vector, non-interacting protein) showed no growth.

(D) AvrPigm promotes the cytoplasmic translocation of HPP04. N-terminal GFP-tagged HPP04 (GFP-HPP04) was transiently expressed in *N. benthamiana* leaves by agroinfiltration together with the indicated proteins. Scale bar, 10 μm.

(E) Analysis of single-gene knockout (KO) mutation sites in HPP04, HIPP16, HIPP19, and HIPP20 in NIL-*Pigm* plants.

(F) Double, triple and quadruple KO mutation sites analysis of HPP04, HIPP16, HIPP19, and HIPP20 in NIL-*Pigm* plants.

(G) Single, Double, and Triple HMA knockout lines in NIL-*Pigm* retain resistant to AvrPigm-containing *M. oryzae*. All the genetic lines listed in the picture at 7dpi with punch injection inoculation (TH12). NIPB served as a susceptible control. Different letters indicate significant differences at $P < 0.05$ ($n = 10$, one way-ANOVA).

(H) Western-blotting showing the expression levels of AvrPigm in NIPB, NIL-*Pigm*, and quadruple *HMA*-KO/ NIL-*Pigm* protoplasts.

(I) High expression of HPP04, HIPP16, HIPP19, and HIPP20 in NIL-*Pigm* caused prevalent lethality in calli and seedlings stage due to autoimmunity. Note that the control calli (expressing corresponding proteins in NIPB) developed greening buds.

(J) Overexpression of HIPP19 caused stunted growth in NIL-*Pigm*.

(K) Overexpression of HPP04 and HIPP19 caused stunted growth in *Piz-t* (ZH11). Left, morphological phenotype of *Ubi::HPP04* and *Ubi::HIPP19* transgenic seedlings in Piz-t; Right, relative expression levels of indicated genes.

(L) Morphology of individual HMA overexpression lines in NIPB. None of the four HMA-overexpressing lines exhibit autoimmunity phenotype. Top, morphological phenotype of HMA overexpression lines in NIPB; Bottom, relative expression levels of the corresponding HMA genes. Scale bar, 10 cm.

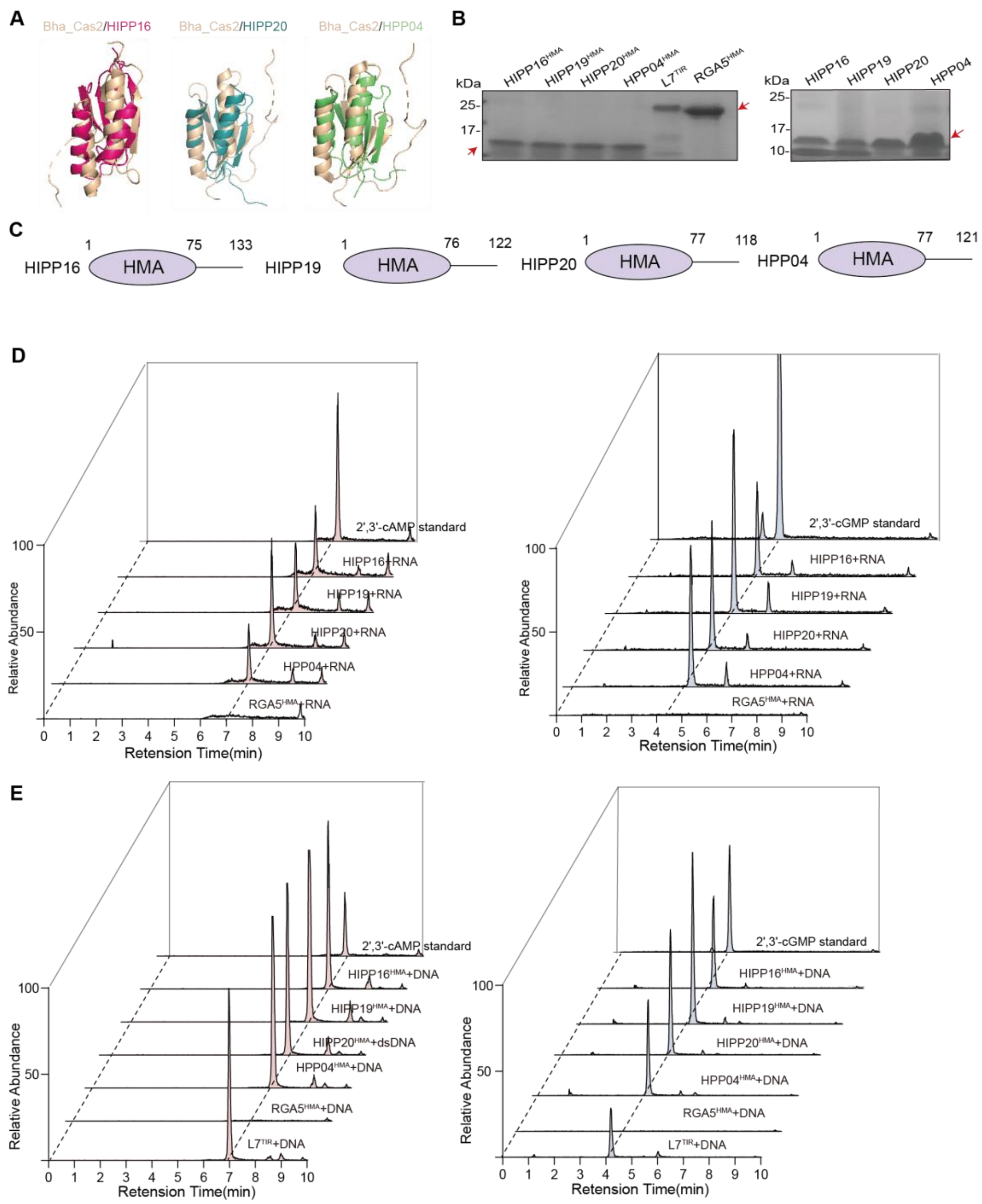


**Figure S3. Four HMA proteins produce 2′,3′-cAMP/cGMP using DNA or RNA as substrates, related to Figure 3.**

(A) Structural alignment of AF3-predicted HMA proteins with Cas2-like nucleases. Structure-based alignment revealed that the monomer structure of Bha_Cas2 (PDB: 4esi, beige) with other predicted HMA-only proteins, including HIPP16$^{HMA}$ (magenta; RMSD = 3.568 Å), HIPP20$^{HMA}$ (teal; RMSD = 4.382 Å) and HPP04$^{HMA}$ (light green; RMSD = 4.913 Å). Structural alignments were performed using PyMOL, with water molecules and metal ions were hided.

(B) SDS-PAGE analysis of purified HMA proteins. His-SUMO2 tagged HIPP16, HIPP19, HIPP20, HPP04, RGA5$^{HMA}$ and His-MBP tagged L7 were purified from *E.coli*, and tags were removed by proteases. Arrows indicated the target tag-free proteins.

(C) Schematic diagram of the domain organization of the four HMA proteins.

(D) LC-MS traces of 2′,3′-cAMP/cGMP production from reactions of four full-length HMA and $RGA5^{HMA}$ proteins incubated with rice RNA. Five HMA proteins were incubated with rice RNA at 25 ℃ for 6 h, and then reaction products were analyzed by LC-MS.(E) LC-MS traces of 2′,3′-cAMP/cGMP from reaction products of four HMA, $RGA5^{HMA}$ and $L7^{TIR}$ proteins incubated with rice genomic DNA. Five HMA proteins and the positive control $L7^{TIR}$ were incubated with rice genomic DNA at 25 ℃ for 16 h, and then reaction products were analyzed by LC-MS relative to their standards. A representative chromatogram is shown for 2′,3′-cAMP (Left) and 2′,3′-cGMP (Right).

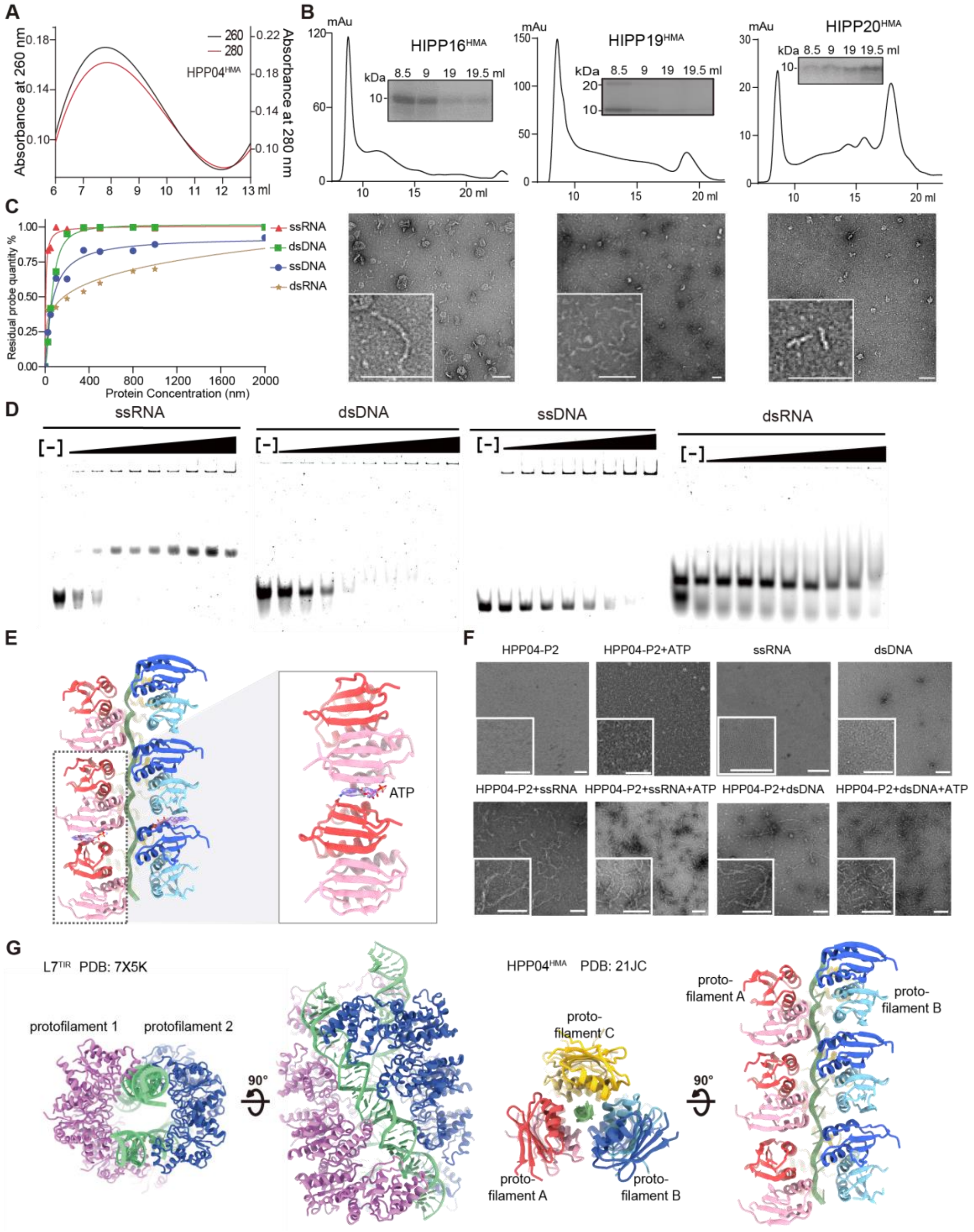


**Figure S4. Biochemical properties and substrate specificity of HMA proteins, related to Figure 4.**

(A) UV absorbance profiles of HPP04$^{HMA}$ filaments. Elution peaks of HPP04$^{HMA}$ filament-forming fractions exhibit high absorbance at 260 nm (A260, black line) and 280 nm (A280, red line), indicating the presence of nucleic acids and protein, respectively.

(B) HIPP16$^{HMA}$, HIPP19$^{HMA}$ and HIPP20$^{HMA}$ form filaments in vitro. SUMO2-tagged HIPP16$^{HMA}$, HIPP19$^{HMA}$ and HIPP20$^{HMA}$ after affinity purification were subjected to a Superose 6 gel filtration column. Fractions eluted at the void position (9 ml) revealed

filament or filament-like structures by negative staining EM as shown on the bottom side. Scale bar, 100 nm.
(C) HPP04$^{HMA}$ preferentially binds ssRNA. Quantification of Electrophoretic mobility shift assay (EMSA) measures HPP04$^{HMA}$ binding affinity to dsDNA, dsRNA, ssDNA, and ssRNA. While HPP04$^{HMA}$ could bind with the above tested nucleic acids, it displays the highest affinity towards ssRNA. Data points are represented by distinct symbols for each nucleic acid type, with lines illustrating binding trends.
(D) The EMSA gel electrophoresis results corresponding to the protein-nucleic acid complexes formed at various nucleic acid concentrations. Each lane represents a different concentration of nucleic acids, and the bands indicate the presence and relative amount of complexes formed.
(E) Cryo-EM density of ATP within the HPP04$^{HMA}$-ssDNA/RNA filament.
(F) Effect of ATP in HPP04$^{HMA}$ filament assembly. Scale bar, 100 nm.
(G) Comparison of HPP04$^{HMA}$ filaments and L7$^{TIR}$ filaments. Differences between the HPP04$^{HMA}$ filament and the L7$^{TIR}$ filament lie in their nucleic-acid composition and the mode of protein-nucleic acid interaction.

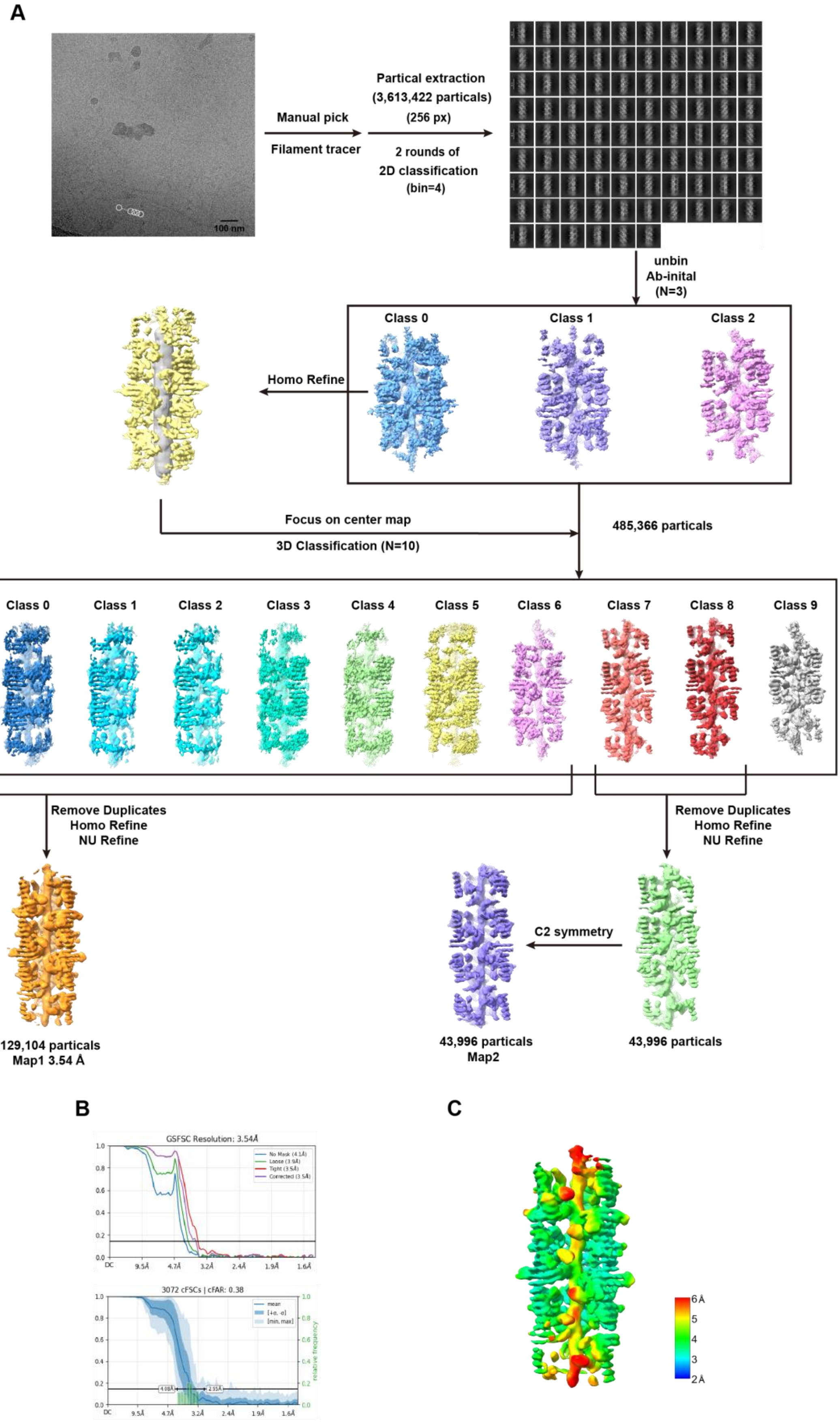
A
100 nm
Manual pick
Filament tracer
Partical extraction
(3,613,422 particals)
(256 px)
2 rounds of
2D classification
(bin=4)
unbin
Ab-inital
(N=3)
Class 0
Class 1
Class 2
Homo Refine
Focus on center map
3D Classification (N=10)
485,366 particals
Class 0
Class 1
Class 2
Class 3
Class 4
Class 5
Class 6
Class 7
Class 8
Class 9
Remove Duplicates
Homo Refine
NU Refine
Remove Duplicates
Homo Refine
NU Refine
C2 symmetry
129,104 particals
Map1 3.54 Å
43,996 particals
Map2
43,996 particals
B
GSFSC Resolution: 3.54Å
3072 cFSCs | cFAR: 0.38
C
6 Å
5
4
3
2 Å

**Figure S5. The 3D reconstruction of HPP04$^{HMA}$ in the complex with ssRNA/DNA, related to Figure 4.**

(A) The flowchart of data processing for the HPP04$^{HMA}$-nucleic acid aggregates complex.

(B) The gold-standard FSC curves of the HMA-nucleic acid aggregates.

(C) The cryo-EM map of the HPP04$^{HMA}$-nucleic acid aggregates, colored by local resolution.

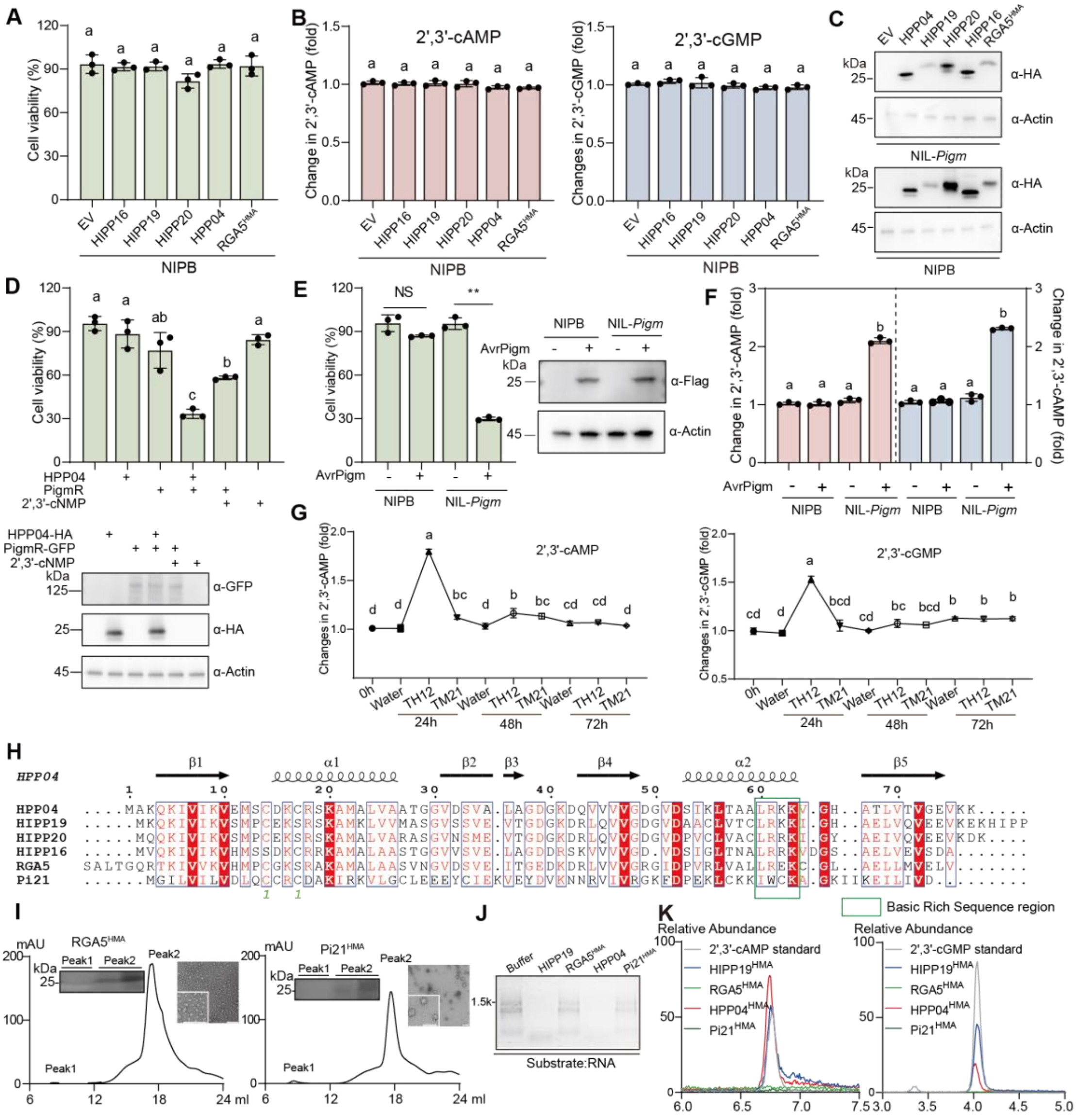

**Figure S6. Overexpression of HMA proteins triggered cell death in NIL-*Pigm* via 2′,3′-cNMP production, related to Figure 5.**

(A) Overexpression of four HMA proteins fail to induce cell death in NIPB protoplasts. The assays were performed by replacing NIL-*Pigm* protoplasts with NIPB protoplasts (Fig. 5A).

(B) 2′,3′-cNMP levels expressing indicated HMA proteins in NIPB. The assays were performed as described for Fig.5B.

(C) Protein levels of HMA proteins in NIL-*Pigm* and NIPB. Western blot analysis of HMA proteins (HPP04, HIPP16, HIPP19, HIPP20, RGA5$^{HMA}$) in NIL-*Pigm* and NIPB protoplasts, detected by anti-HA with anti-Actin as a loading control.
(D) 2′,3′-cNMP triggers cell death in NIPB protoplasts expressing PigmR. Nipponbare protoplasts expressing PigmR exhibit reduced cell viability after treatment with 1 mM 2′,3′-cNMP. Protein levels were detected for indicated constructs.
(E) AvrPigm triggers cell death in NIL-*Pigm* but not in NIPB. Protoplasts expressing AvrPigm or empty vector and cell viability was measured using Cell Titer-Glo Luminescent Assay after 12 h post-transfection. Protein levels of AvrPigm were detected in NIL-*Pigm* and NIPB
(F) Expression of AvrPigm elevates 2′,3′-cNMP levels in NIL-*Pigm* protoplasts. Identical workflow to Fig 5B.
(G) 2′,3′-cNMP levels in compatible (TM21) and incompatible (TH12) interactions between *M. oryzae* and NIL-*Pigm*. Water treatment serves as control.
(H) Sequence alignment of selected rice HMA proteins. Green square indicates BRS region in α helix, sequence alignment was performed by MUSCLE.
(I) Gel filtration and negative-staining images of some HMA proteins eluted at the void position from Superose 6 gel filtration column. RGA5$^{HMA}$ and Pi21$^{HMA}$ proteins lack the BRS region and fail to form filaments.
(J, K) Nuclease (J) and 2′,3′-cNMP synthetase (K) activities of indicated HMA proteins towards RNA. LC-MS trace for 2′,3′-cNMP standard and the products that HMA proteins catalyzed RNA. The assays for the nuclease and 2′,3′-cNMP synthetase activities were performed (Fig 3C and 3F). RGA5$^{HMA}$ and Pi21$^{HMA}$ proteins do not possess nuclease and 2′,3′-cNMP synthetase activities.
Asterisks represent significant differences (*$P < 0.05$, **$P < 0.01$) in E. Different letters indicate significant differences at $P < 0.05$ (n=3, one way-ANOVA) in A, B, D, F, G.

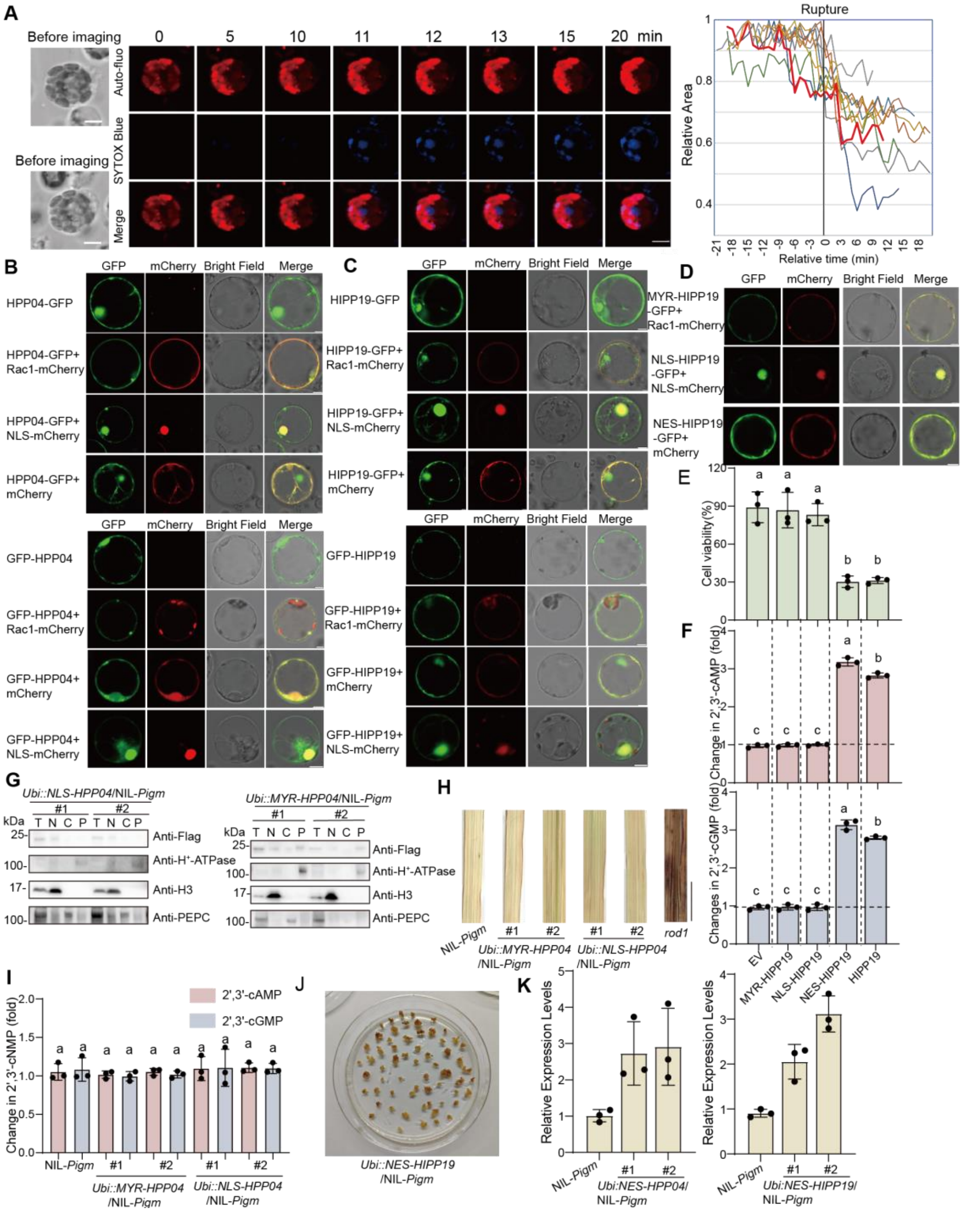

**Figure S7 Cytoplasmic HMA proteins induced PigmR-mediated cell death, related to Figure 6.**

(A) Time-lapse imaging of PigmR-triggered cell death in protoplasts. NIL-*Pigm* protoplasts expressing HPP04 were stained with SYTOX Blue and monitored by time-lapse microscopy to visualize cell death process (n =10 cells).

(B) Subcellular localization of HPP04/HIPP19-GFP in rice protoplasts. Protoplasts expressing HPP04/HIPP19-GFP with subcellular markers: plasma membrane (Rac1-mCherry, magenta), nucleus (NLS-mCherry, magenta), or cytoplasm (mCherry, magenta). Merged images demonstrate co-localization. Samples were visualized under a confocal microscope, individual fluorescence channels, merged images, and bright

field images are shown.
(C) Subcellular localization of GFP-HPP04 and GFP-HIPP19 in rice protoplasts. Identical workflow to (C), replacing C-terminal GFP with N-terminal GFP of HPP04 and HIPP19.
(D) Subcellular localization of MYR/NLS/NES-HIPP19-GFP in NIL-*Pigm* protoplasts. Identical workflow to Fig.6F, replacing HPP04 with HIPP19.
(E, F) Subcellular localization modulates HIPP19-induced cell death (E) and 2′,3′-cNMP levels in NIL-*Pigm*. Identical workflow to Fig.6G,6H, replacing HPP04 with HIPP19.
(G) Subcellular fraction of proteins from NLS/MYR-HPP04 in NIL-*Pigm* plants. Total protein (T) was extracted and separated into nuclear (T), cytoplasm (C) and PM (P) fractions by using the Minute PM protein isolation kit. Immunoblotting with α-Flag, α-$H^+$-ATPase (PM), α-H3 (nucleus) and α-PEPC (cytoplasm).
(H) DAB staining for $H_2O_2$ accumulation in *NLS-HPP04*/NIL-*Pigm* and *MYR-HPP04*/NIL-*Pigm* transgenic lines. Scale bar, 1 cm.
(I) The 2′,3′-cNMP levels in *NLS-HPP04*/NIL-*Pigm* and *MYR-HPP04*/NIL-*Pigm* transgenic lines are not significantly different from those in NIL-*Pigm*. Statistical analysis of 2′,3′-cAMP/cGMP levels are performed independently.
(J) Cell death of transgenic rice calli expressing NES-HIPP19 in NIL-*Pigm*.
(K) Relative expression levels of HIPP19 and HPP04 in *NES-HIPP19* or *HPP04* in NIL-*Pigm*, as detected by qRT-PCR. Each point represents an independent biological line.
Scale bar, 5 μm in A, B, C, D. Different letters indicate significant differences at $P < 0.05$ (n=3, one way-ANOVA) in E, F, I.

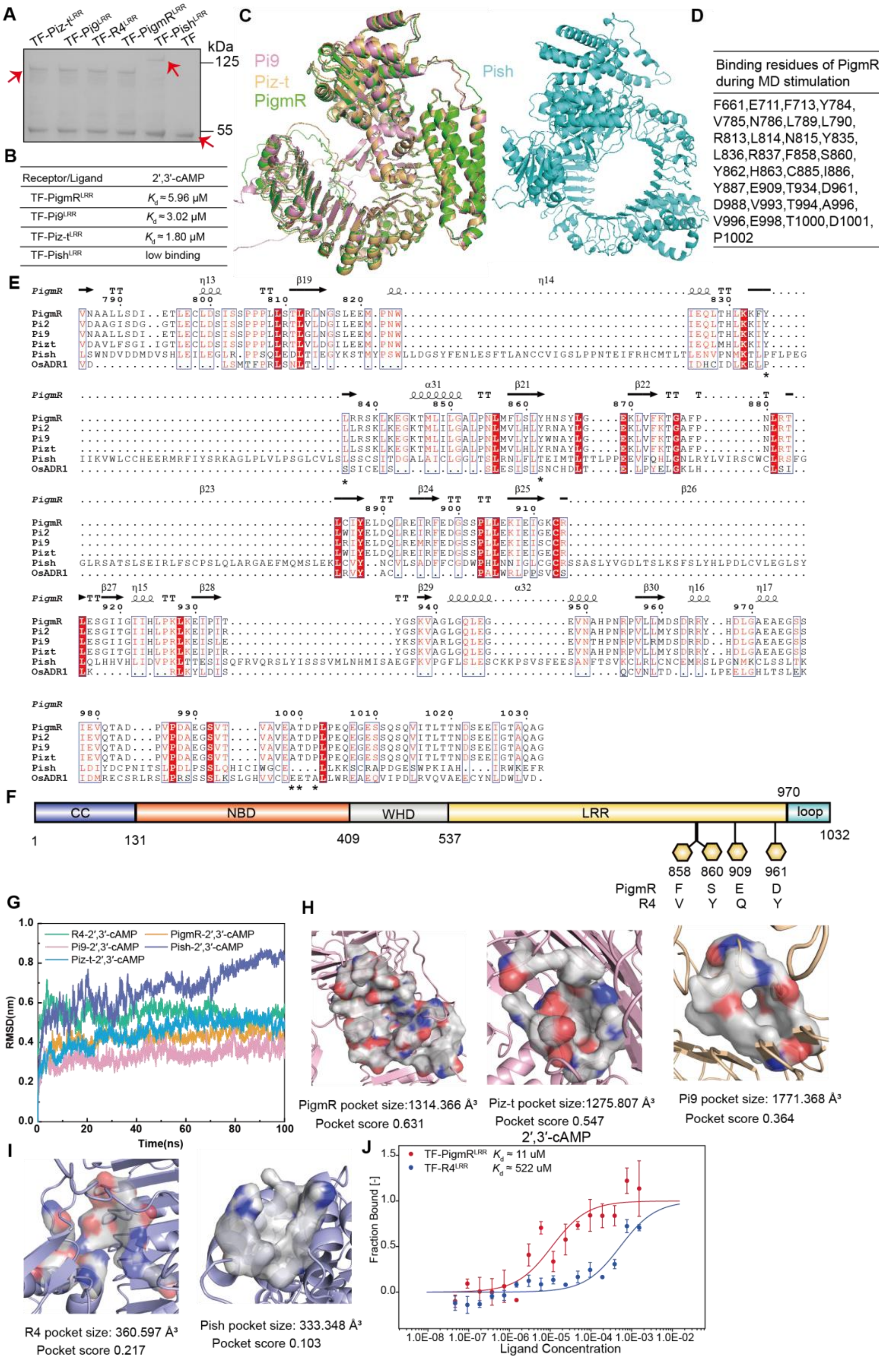


**Figure S8. The LRR domains of NLRs mediates the recognition of 2′,3′-cNMP, related to Figure 7.**

(A) SDS-Page analysis of purified the LRR domains of NLRs (Arrowheads).
(B) Quantification of the binding affinity between PigmR/Piz-t/Pi9/Pish LRR domains and 2′,3′-cAMP, measured by FIDA.
(C) The predicted protein structure of Pi9, Piz-t, PigmR and Pish through AF3. pTM score of NLRs: PigmR (0.82), Pi9 (0.84), Piz-t (0.83) and Pish (0.77). A pTM score above 0.5 means the overall predicted fold for the complex might be similar to the true structure.
(D) Identified critical residues for PigmR-2′,3′-cAMP during molecular dynamics (MD) stimulation.
(E) Sequence alignment of LRR domains from multiple NLRs. PigmR, Pi2, Pi9, and Piz-t are paralogs from the same gene cluster, whereas Pish and OsADR1 are encoded on different chromosomes.
(F) Domain architecture of PigmR. Schematic representation of the functional domains of PigmR. SNPs (hexagon) distinguishes PigmR from non-functional paralogous NLR, R4.
(G) Root mean square deviation (RMSD) of 2′,3′-cNMP in the interaction modes with multiple NLRs, generated by MD simulation. The average RMSD values for the R4-2′,3′-cAMP, PigmR-2′,3′-cAMP, Pi9-2′,3′-cAMP, Pish-2′,3′-cAMP and Piz-t-2′,3′-cAMP systems were 0.541 ±0.038 nm, 0.420 ±0.047 nm, 0.333±0.042 nm, 0.669±0.11 nm and 0.45±0.07 nm, respectively. The smaller fluctuation amplitudes indicate higher structural stability during the simulation, demonstrating good reliability of the systems.
(H) Binding pocket analysis of PigmR and other NLRs during MD stimulation. PigmR and its functional paralog NLRs (Pi9 and Piz-t) demonstrated enlarged ligand-binding pocket volumes with high pocket scores.
(I) Binding pocket analysis of R4 and Pish during MD stimulation. R4 and Pish displayed smaller ligand-binding pocket volumes with lower pocket scores.
(J) Quantification of binding between the LRR domains of PigmR/R4 and 2′,3′-cAMP by Microscale Thermophoresis (MST).
